# Modulation of motor behavior by the mesencephalic locomotor region

**DOI:** 10.1101/2020.06.25.172296

**Authors:** Daniel Dautan, Adrienn Kovács, Tsogbadrakh Bayasgalan, Miguel A. Diaz-Acevedo, Balazs Pal, Juan Mena-Segovia

## Abstract

The mesencephalic locomotor region (MLR) serves as an interface between higher-order motor systems and lower motor neurons. The excitatory module of the MLR is composed of the pedunculopontine nucleus (PPN) and the cuneiform nucleus (CnF), and their activation has been proposed to elicit different modalities of movement, but how the differences in connectivity and physiological properties explain their contributions to motor activity is not known. Here we report that CnF glutamatergic neurons are electrophysiologically homogeneous and have short-range axonal projections, whereas PPN glutamatergic neurons are heterogeneous and maintain long-range connections, most notably with the basal ganglia. Optogenetic activation of CnF neurons produced fast-onset, involuntary motor activity mediated by short-lasting muscle activation. In contrast, activation of PPN neurons produced long-lasting increases in muscle tone that reduced motor activity and disrupted gait. Our results thus reveal a differential contribution to motor behavior by the structures that compose the MLR.

## Introduction

The mesencephalic locomotor region (MLR) is a functionally-defined midbrain area composed of the pedunculopontine nucleus (PPN) and the cuneiform nucleus (CnF) which has been typically described as an output station of forebrain systems reaching lower motor circuits ^1–3^. Early experiments defined the MLR by demonstrating that electrical stimulation of this region induced a locomotor response in decorticated cats^4,5^. More recently, optogenetic experiments revealed that the motor function of the MLR is dependent on excitatory transmission from glutamatergic neurons^6,7^, which is the most prominent cell type in the MLR^8,9^. In the last two decades, a role for these circuits in gait and posture has been proposed^10–13^. Moreover, degeneration of neurons in the MLR may underlie some of the motor impairments in Parkinson’s disease^14–19^. Deep brain stimulation into the PPN has been shown to produce some improvements in abnormal gait based on the idea that the output from the MLR is excitatory^20–23^. However, it is not fully understood how excitatory MLR neurons contribute to motor behavior and how motor functions are associated with different neuronal types in the MLR.

The PPN, the largest component of the MLR, is highly heterogeneous. It is composed of three neurotransmitter-defined cell types: cholinergic, GABAergic and glutamatergic neurons. Among PPN glutamatergic neurons, a high degree of variability has been reported in their neurochemical composition^8^, connectivity^24^ and firing properties^25^. Comparatively less is known about the CnF. Recent reports show that activation of CnF glutamatergic neurons induces a robust motor activation that is functionally distinct from the activation of PPN neurons, suggesting a functional specialization of MLR neurons^7,12^. PPN and CnF are contiguous structures, the borders of which are not well defined^26^, and this imposes a challenge for unambiguously separating both populations. Furthermore, there is a certain level of interconnectivity that accounts for an additional degree of difficulty in the interpretation of functional studies. We, therefore, sought to identify the functional properties of PPN and CnF neurons and their involvement in motor control.

We used a range of electrophysiological, behavioral and anatomical techniques to dissect the properties of glutamatergic neurons in the PPN and CnF and identify their specific contributions to motor function and muscle activity. Our results establish fundamental differences in the properties and functions of the PPN and CnF.

## Results

### Input/output connectivity of PPN and CnF with segregated motor circuits

To determine the afferent and efferent connectivity of MLR glutamatergic neurons, we used Cre-dependent anterograde and retrograde viral tracing strategies in VGLUT2-Cre mice. Given the proximity of MLR structures, only microinjections that were strictly confined to the borders of the PPN or the CnF were considered further. We used the immunolabeling of choline acetyltransferase (ChAT) to delimit the boundaries of the PPN and the ventral border of the CnF^26–29^. CnF is ventrally bordered by the PPN and dorsally by the central nucleus of the inferior colliculus and fibrodendritic lamina^30^. A glutamatergic neuron was considered to belong to the PPN if it was located within 100μm of the closest cholinergic neurons (**Fig. 1A**), or to the CnF if it was located at least 100μm dorsally to cholinergic neurons and ventral to the cIC (**Fig. 1B**). If more than 5% of virus-labeled neurons were located outside the borders of the targeted structure, the data from the animal were excluded (**Supplementary Fig. 1A**).

**Figure 1.**
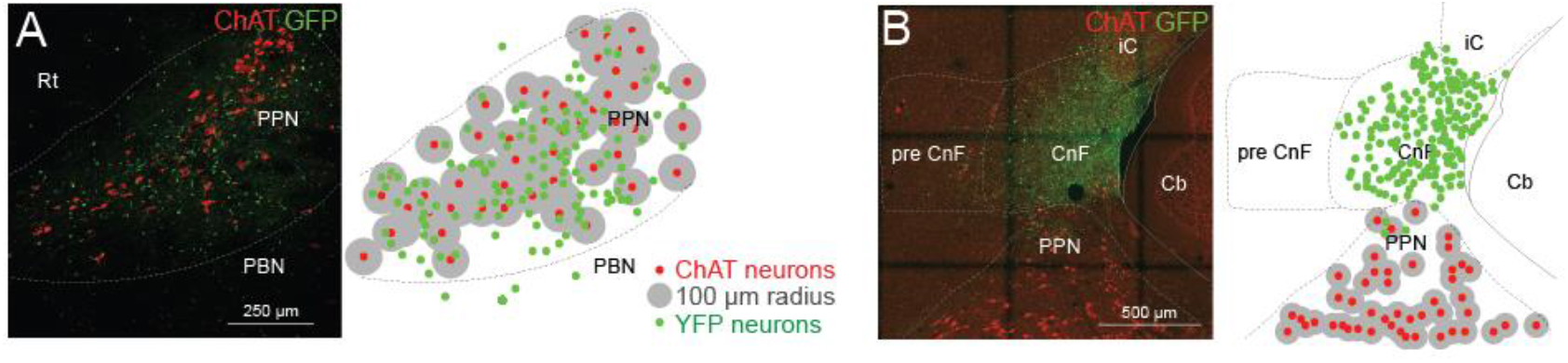
Segregation of MLR structures by viral transduction. Viral injection volume was adjusted to be restricted within the border of the PPN (**A**) or the CnF (**B**) using as a marker the expression of choline acetyltransferase (ChAT; 100μm radius). The dorsal and ventral border of the PPN was defined as 100μm distance from the outer cholinergic neuron soma (**A**), whereas the ventral border of CnF was 100μm further from cholinergic neurons (**B**).

To label the axonal projections and synapses originating in the PPN and CnF, we transduced glutamatergic neurons with a reporter expressing cytosolic green fluorescent protein (GFP) in the presence of Cre recombinase and a red fluorescent protein (mRuby) under the control of the promoter for the pre-synaptic marker synaptophysin (AAV-hSyn-FLEX-GFP-2A-Synaptophysin-mRuby, PPN n = 3; **Fig. 2A1**; CnF n = 3; **Fig. 2A2**). We next mapped the synaptophysin expression across the brain and spinal cord using high-resolution confocal imaging (**Fig. 2B, F**). No differences in the number of transduced neurons (GFP^+^) between PPN and CnF were observed (**Fig. 2C, Supplementary Fig. 1B-C**). However, the density of synapses (mRuby^+^/GFP^+^ puncta) was higher in the PPN group compared to the CnF group (**Fig. 2D**). We next compared the distribution of synapses across the brain between both groups (**Fig. 2B, E**). We found comparatively more innervation by PPN than CnF neurons in the basal ganglia (PPN: 0.336 ± 0.089 pixels/100μm^2^; CnF: 0.095 ± 0.032 pixels/100μm^2^; Wilcoxon rank-sum test *P* = 0.0369; **Fig. 2F**), as well as in individual structures such as the dorsal raphe (PPN: 0.56 ± 0.121 pixels/100μm^2^; CnF: 0.047 ± 0.023 pixels/100μm^2^; *P* = 0.0292) and the dorsal periaqueductal gray area (dPAG; PPN: 0.251 ± 0.078 pixels/100μm^2^; CnF: 0.176 ± 0.034 pixels/100μm^2^; *P* = 0.037), in agreement with previous tracing studies^31–33^. In contrast, the innervation originated in CnF glutamatergic neurons was mostly concentrated in the midbrain and similar to the PPN (PPN: 0.25 ± 0.028 pixels/100μm^2^; CnF: 0.17 ± 0.029 pixels/100μm^2^; *P* = 0.0562) and the pons (PPN: 0.14 ± 0.03 pixels/100μm^2^; CnF: 0.12 ± 0.033 pixels/100μm^2^; *P* = 0.56; **Fig. 2G**), including the tectal area, the parabrachial nucleus, PAG, and the ventral gigantocellular nucleus, in agreement with previous studies^34–36^. Furthermore, PPN but not CnF neurons show synaptic labeling in the striatum, the substantia nigra pars compacta (4.27 ± 0.12 pixels/100μm^2^), cerebellum (4.19 ± 0.04 pixels/100μm^2^), the dorsal brainstem (vestibular nucleus: PPN 0.77 ± 0.02 pixels/100μm^2^, CnF: 0 pixels/100 μm^2^; medullary reticular nucleus, MdV: PPN 0.4 ± 0.01 pixels/100 μm^2^, CnF: 0 pixels/100 μm^2^; **Fig. 2G**) and the spinal cord (**Fig. 2J**). PPN glutamatergic projections to the spinal cord were observed at all segments (**Fig. 2K**), but the synaptic density was not quantified. Following axonal reconstructions of randomly selected cervical, thoracic, lumbar and sacral segment sections (**Fig. 2K**), we found that PPN projections follow the rubrospinal tract and decussate in laminae 5-7, making dense synapses (mRuby signal) that seem to avoid motor neurons, as revealed with ChAT-immunostaining, which is expressed in spinal neurons. Taken together, these findings suggest that PPN directly projects to structures involved in different modalities of movement (i.e. basal ganglia, brainstem, cerebellum and spinal cord), whereas CnF sends projections to structures involved in the execution of movement (i.e. ventral gigantocellular nucleus; **Supplementary Fig. 2**).

**Figure 2.**
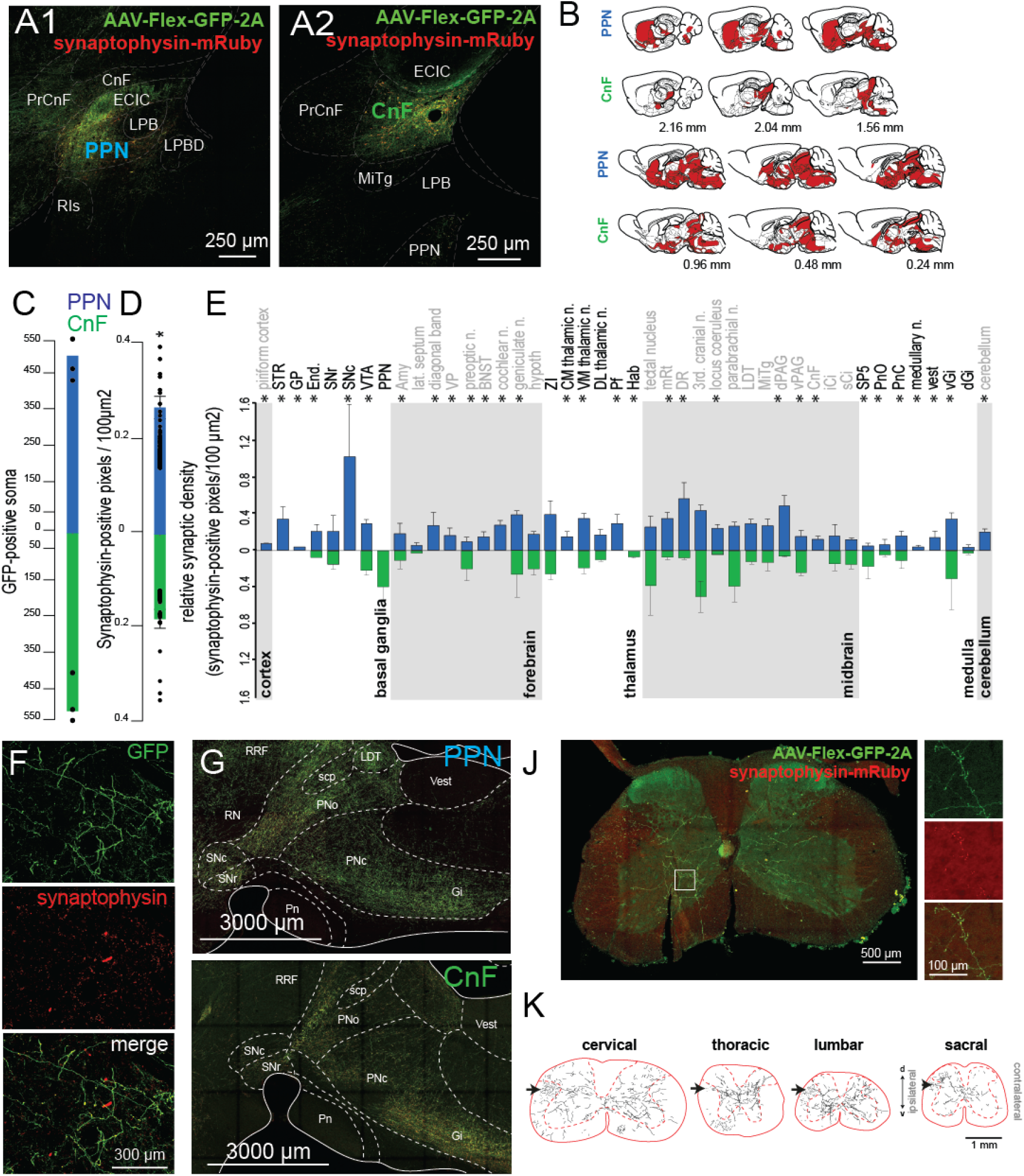
Axonal distribution of PPN and CnF glutamatergic neurons. **A-B,** Following injection of AAV-DIO-GFP-2A-synaptophysin-mRuby (**A**) restrained to the PPN (**A1**) or the CnF (**A2**) borders, we observed widespread distribution of GFP-labeled axons (**B**). **C-D,** Quantification of the total count of GFP-positive soma (PPN: 504.66 ± 58.42; CnF: 518.33 ± 54.67, one-way ANOVA F_(1,5)_ = 0.03, *P* = 0.87) and overall synaptic density (PPN: 0.27 ± 0.024 pixels/100 μm^2^; CnF: 0.18 ± 0.02 pixels/100 μm^2^, one-way ANOVA F_(1,86)_ = 7.64, *P* = 0.007). **E,** Segregated synaptophysin labeling across the brain revealed distinct patterns of innervation by PPN and CnF glutamatergic neurons, particularly in the basal ganglia, forebrain, thalamus, midbrain, medulla and cerebellum (Wilcoxon test). **F,** Fluorescent micrographs illustrating GFP and synaptophysin labeling in the striatum following PPN transduction. **G,** Distribution of axons in the brainstem following PPN and CnF injections. **J,** Synaptic distribution in a cervical segment of the spinal cord. **K**, Axonal reconstructions in typical examples of cervical, thoracic, lumbar, and sacral spinal cord segments following unilateral PPN injection. *P<0.05. All experiments have been replicated at least 3 times. Single data are represented by small dots. All data are represented as mean ± SEM.

Next, to identify the inputs to the glutamatergic neurons of the PPN and CnF, we used a monosynaptic retrograde labeling strategy in VGLUT2-cre mice (RvDG-YFP). Because the specificity of the retrograde tracing is conferred by the expression of the helper viruses, we adjusted the volume of the helper mix (AAV-DIO-TVA-mCherry, AAV-DIO-Rg, 1:1) to selectively target PPN (**Fig. 3A**; n=3) or CnF (**Fig. 3B**; n=3) according to the criteria described earlier. Starter neurons (mCherry-/YFP-positive) within the PPN and CnF were located within the border of each structure (**Fig. 3C**). The overall number of input neurons (YFP-positive) was larger in the PPN group compared to the CnF group (**Fig. 3D**). After normalization by the number of starter neurons, the number of input neurons was still larger in the PPN group (PPN: 4.73+/−1.27, CnF: 2.28+/−0.94, P=0.027). We found a larger number of input neurons to the CnF originating in the colliculi (PPN: 22 ± 1.74, CnF: 39.66 ± 8.46, Wilcoxon rank-sum *P* = 0.0033; **Fig. 3E**), the PAG (PPN: 5.06 ± 0.44, CnF : 8.52 ± 0.92, *P* = 0.0275) and the precuneus (PPN: 7.46 ± 0.61, CnF: 15.238 ± 1.89, *P* = 0.0175), whereas a larger number of input neurons to the PPN originated in the PnO (PPN: 7.47 ± 0.21, CnF: 2.21 ± 1.46, *P* = 0.0234, **Fig. 3H**), the striatum (PPN: 2.66 ± 1.55, CnF : 0.33±1.15, *P* = 0.04, **Fig. 3I**), the ZI (PPN: 4.33 ± 0.15, CnF: 1.18 ± 0.70, *P* = 0.025) and the motor cortex (PPN: 4.66 ± 0.66, CnF : 1.52 ±1.52, *P* = 0.025; **Fig. 3J**). Input neurons in a subset of motor structures were observed to connect exclusively with the PPN, including the SNr (PPN: 5.3 ± 0.39, CnF : 0; **Fig. 3F-G**), spinal cord (PPN: 7.23 ± 0.96, CnF : 0; **Fig. 3K**), gigantocellular nucleus (PPN: 0.98 ± 0.22, CnF: 0), dorsal gigantocellular nucleus (PPN: 2.16 ± 0.20, CnF: 0), paragigantocellular nucleus (PPN: 1.33 ± 0.13, CnF: 0) and deep cerebellar nuclei (PPN: 4.22 ± 0.55, CnF: 0; **Figure 3E-K, Supplementary Fig. 2**). Overall, the distribution of inputs to PPN glutamatergic neurons is far more widespread than the distribution of inputs to CnF glutamatergic neurons and largely overlaps with the PPN/CnF output targets (**Figure 3L**). Combined, these results reveal differences in the input/output connectivity of PPN and CnF glutamatergic neurons with separate motor circuits (**Supplementary Figure 2**).

**Figure 3.**
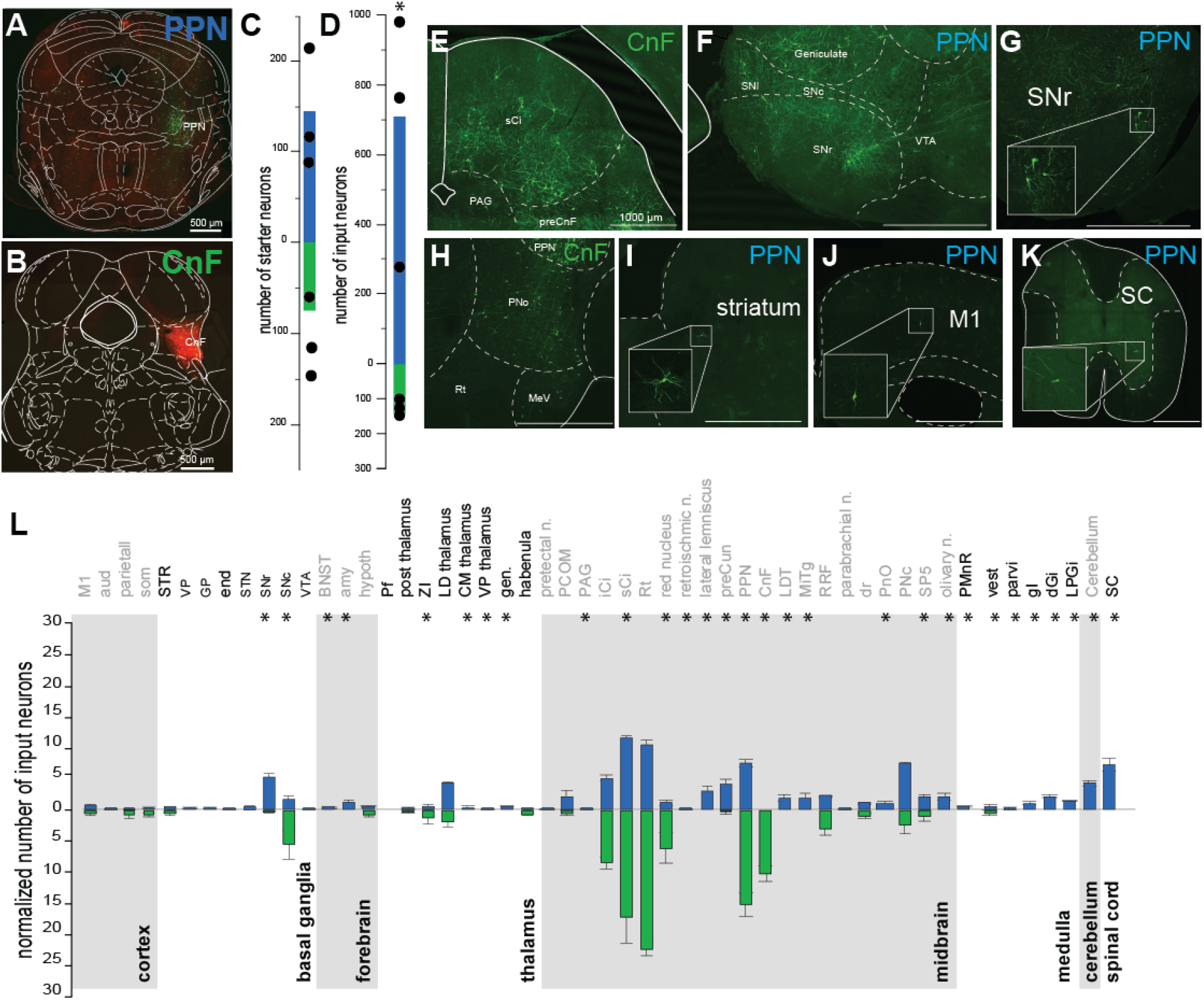
Whole-brain inputs to PPN and CnF glutamatergic neurons. **A-D,** Following injection of helpers and rabies virus in the PPN (**A**) and the CnF (**B**), we quantified the number of starter neurons (**C**, PPN: 145.33 ± 40.19, CnF: 74.66 ± 24.91, t-test t_(4)_ = 1.49, *P* = 0.21047) and the number of inputs neurons across all brain areas (**D**, raw: PPN: 708.91 ± 242.25, CnF: 143.33 ± 12.17, t-test t_(4)_ = 2.33, *P* = 0.0401; normalized: PPN: 4.73 ± 1.27, CnF: 2.28 ± 0.94 input/starter, Mann-Whitney: Z = 1.964, *P* = 0.0495). **E-K,** Fluorescent micrographs of representative areas where inputs neurons were identified, including the dorsal brainstem (**E**) and the pons (**H**) following a CnF injection, and the ventral midbrain (**F-G**), the striatum (**I**), the cortex (**J**) and the spinal cord (**K**) following a PPN injection. **L,** Quantification of the number of inputs neurons projecting to PPN (blue) and CnF(green) glutamatergic neurons for each brain area normalized by the overall total number of input neurons per animal (Wilcoxon test). *P<0.05. All experiments have been replicated at least 3 times. Single data are represented by small dots. All data are represented as mean ± SEM.

### Glutamatergic PPN neurons are physiologically distinct to CnF neurons

To characterize the physiological properties of MLR glutamatergic neurons, we obtained brain slices for *ex vivo* recordings of identified PPN (n=77) and CnF (n=41) glutamatergic neurons of VGLUT2-tdTomato mice (**Figure 4, Supplementary Fig. 3, Supplementary Table 2-3**). From the recorded td-Tomato-positive neurons, randomly selected subsets (PPN n=15, CnF n=11; **Fig. 4A-B**) were subsequently labeled and reconstructed, revealing that CnF glutamatergic neurons have a significantly larger number of main dendrites, nodes and endings (**Fig. 4C-D; Supplementary Table 2**). Based on the classical electrophysiological classification of PPN neurons^37,38^ (**Supplementary Figure 3, Supplementary Tables 2-3**), we defined functional subgroups based on changes of spike frequency adaptation with increasing depolarization^39^ and classified neurons in 3 groups: non-adapting, slowly adapting and rapidly adapting (**Fig. 4E-G**). In the PPN, 30.2% of all neurons (13/43 neurons) were non-adapting and were located predominantly in the lateral regions, whereas 21% (9/43 neurons) were slowly adapting and 48.8% (21/43 neurons) were rapidly adapting. In contrast, in the CnF the large majority of neurons (85.7%, 24/28 neurons) were rapidly adapting, and non-adapting and slowly adapting constituted equal smaller proportions (7.15%, 2/28 neurons for each category; **Fig. 4H**). Thus, the responses of MLR glutamatergic neurons to spike adaptation reveal important biophysical group differences in the composition of the PPN and the CnF ranging from firing frequency to adaptation index (**Supplementary Fig. 4, Supplementary Table 2**).

**Figure 4.**
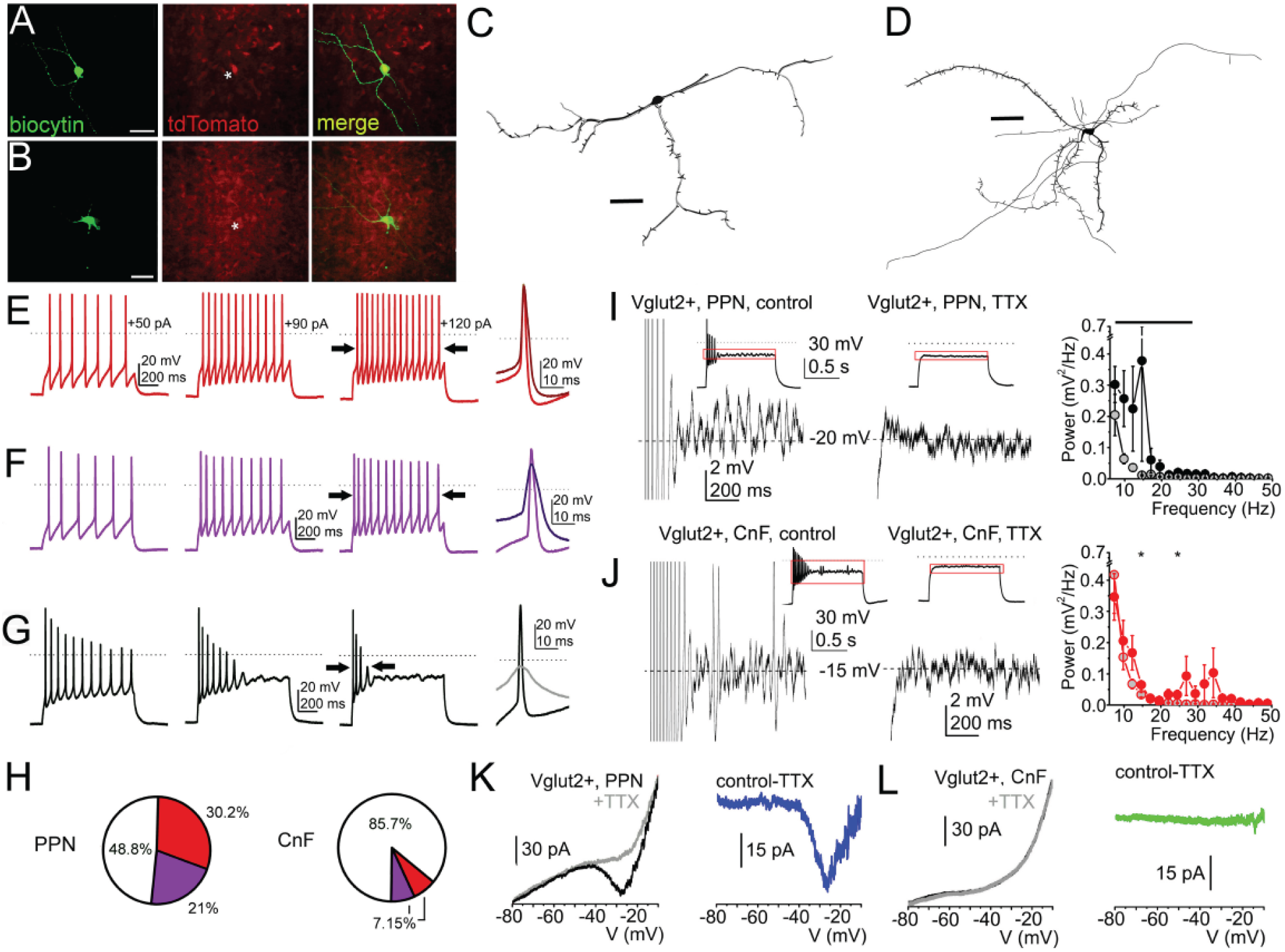
Functional and morphological differences of PPN and CnF glutamatergic neurons. **A-B,** Fluorescent micrographs of PPN and CnF glutamatergic neurons obtained from VGLUT2-tdTomato mice following biocytin labelling. **C-D,** Reconstruction of representative glutamatergic neurons in the PPN (**C**) and the CnF (**D**), which were subsequently used to quantify the number of proximal dendrites, nodes and endings (**Supplementary Table 2**). **E-G,** Changes of spike frequency adaptation by increasing depolarizing steps revealed functional subtypes of glutamatergic neurons defined as follows, (**E**) ‘non-adapting’: less than 50% increase in the adaptation index of the action potential trains obtained with 50 and 120 pA current injections; (**F**) ‘slowly-adapting’: more than 50% change of the adaptation index but fired during the whole 1-s-long depolarizing step; and (**G**) ‘rapidly-adapting’: paused firing after application of greater depolarizing steps. **H,** Proportion of neurons with different spike frequency adaptation properties in the PPN and the CnF. **I-J,** Voltage traces from glutamatergic neurons in the PPN (**I**) and the CnF (**J**) representing high threshold oscillations during 120 pA depolarizing square current injections under control conditions (left) and following TTX application (right; red squares of the small inserts indicate the magnified area). Related power spectra are displayed on the left (average ± SEM; PPN control, black circles; PPN+TTX, gray circles with black contours; CnF control, red circles; CnF+TTX, gray circles with red contours). **K-L,** Representative current traces from neurons in the PPN (**K**) and the CnF (**L**) elicited by voltage ramp injections under control conditions (black) and with TTX (gray; left). TTX-sensitive currents shown on the right panels (PPN, blue; CnF, green). Scale bars: A-B: 0.5mm, C-D: 50μm. * P<0.05. All experiments have been replicated at least 3 times. Group value and statistics are provided in Table 2. All data are represented as mean ± SEM.

Next, because neurons in the MLR region have been reported to display high-threshold membrane potential oscillations^40,41^, we sought to characterize the oscillatory activity of glutamatergic neurons of the PPN and CnF. Oscillatory activity in the 10-20 Hz range was present in PPN glutamatergic neurons (n=24 neurons) and was sensitive to TTX (**Fig. 4I**). In contrast, oscillatory activity in the 20-40 Hz range was present in the CnF (n=19 neurons) but it was weaker and largely insensitive to TTX (**Fig. 4J**). Power spectra revealed similar average frequency ranges of oscillatory activity in the PPN and CnF, but with a greater standard deviation in CnF neurons. TTX-resistant oscillations were virtually absent in both structures (**Supplementary Table 2**). Furthermore, persistent sodium currents were observed predominantly in the PPN (9/11 neurons, range from 26 to 58.7 pA, average 32.5 ± 4.5 pA; **Fig. 4K**), and to a much lesser extent in the CnF (3/7 neurons, range from 7 to 22.2 pA, average 12.4 ± 4.9 pA; **Fig. 4L**), suggesting their likely contribution to the oscillatory activity observed in PPN neurons. In summary, PPN glutamatergic neurons form a heterogeneous group and display robust, wide-range oscillatory activity and the presence of persistent sodium currents. In contrast, CnF glutamatergic neurons are largely fast-adapting and mostly lack persistent sodium currents.

### PPN and CnF activation produces contrasting effects on motor activity

The differences in the connectivity and physiological properties between PPN and CnF reported here suggest that neurons in each structure are recruited by different motor circuits and that their dynamics of activation differ. Recent reports have shown that CnF neurons modulate speed locomotion^6,12,42^, whereas PPN neurons have been suggested to modulate exploratory locomotion^7^ and locomotion pattern^12^. To elucidate the extent of overlap of PPN and CnF function in the context of motor behavior, we used an optogenetic strategy to stimulate glutamatergic neurons while mice were tested in a battery of motor tasks. We unilaterally transduced ChR2 into the PPN or the CnF of VGLUT2-Cre mice and implanted an optic fiber to deliver blue light and activate ChR2 (**Supplementary Fig. 1A)**. First, we determined the effects in the open field (40×40cm, **Fig. 5A**). Stimulation of CnF (n=6 mice), but not PPN (n=8 mice), increased motor activity (CTRL: n=8 mice; **Fig. 5B-E**). The analysis of the individual trials revealed that stimulation of CnF glutamatergic neurons robustly increased the distance traveled compared to the baseline (**Fig. 5B**). In contrast, stimulation of PPN glutamatergic neurons (**Fig. 5C**) significantly reduced the distance traveled. To determine whether the stimulation effects were dependent on the behavioral state of the animal, we separated the stimulation trials based on whether animals were moving or not. We found that CnF stimulation increases the distance traveled and speed regardless of the behavioral state of the animal (**Fig. 5D**), whereas PPN effects were only visible during ongoing movement (**Fig. 5E**). Because the effects of PPN stimulation reported here contrast with previous studies that reported an increase in motor activity during PPN activation^6,7^, we next explored whether different stimulation protocols may account for the differences between studies. We used three stimulation frequencies (1, 10 and 20Hz, 1s ON/9s OFF; **Fig. 5F-G**) and found that, in line with the effects reported above, the effect of PPN stimulation was consistently inhibitory, whereas the increase in motor activity elicited by CnF stimulation was frequency-dependent (**Fig. 5F-G**). No differences in the time spent in center vs periphery of the open field were detected (**Fig. 5H**), thus ruling out an anxiogenic effect of the stimulation. To further characterize the frequency-dependent effects reported above, we tested a different cohort of CnF-transduced and -implanted animals (n=6) in a larger open field (80 x 80cm) with randomized stimulation frequencies ranging from 0.1 Hz to 30 Hz (1s ON/9s OFF). We found that all stimulation frequencies increased the distance traveled (**Supplementary Fig. 5A**) with a maximum effect observed at 12.5Hz and that the effect was restricted to the duration of the stimulation (**Supplementary Fig. 5B, Video 1**). These results reveal that CnF stimulation increases the distance traveled by generating robust and consistent bouts of motor activity, whereas stimulation of PPN neurons briefly reduces exploratory locomotion.

**Figure 5.**
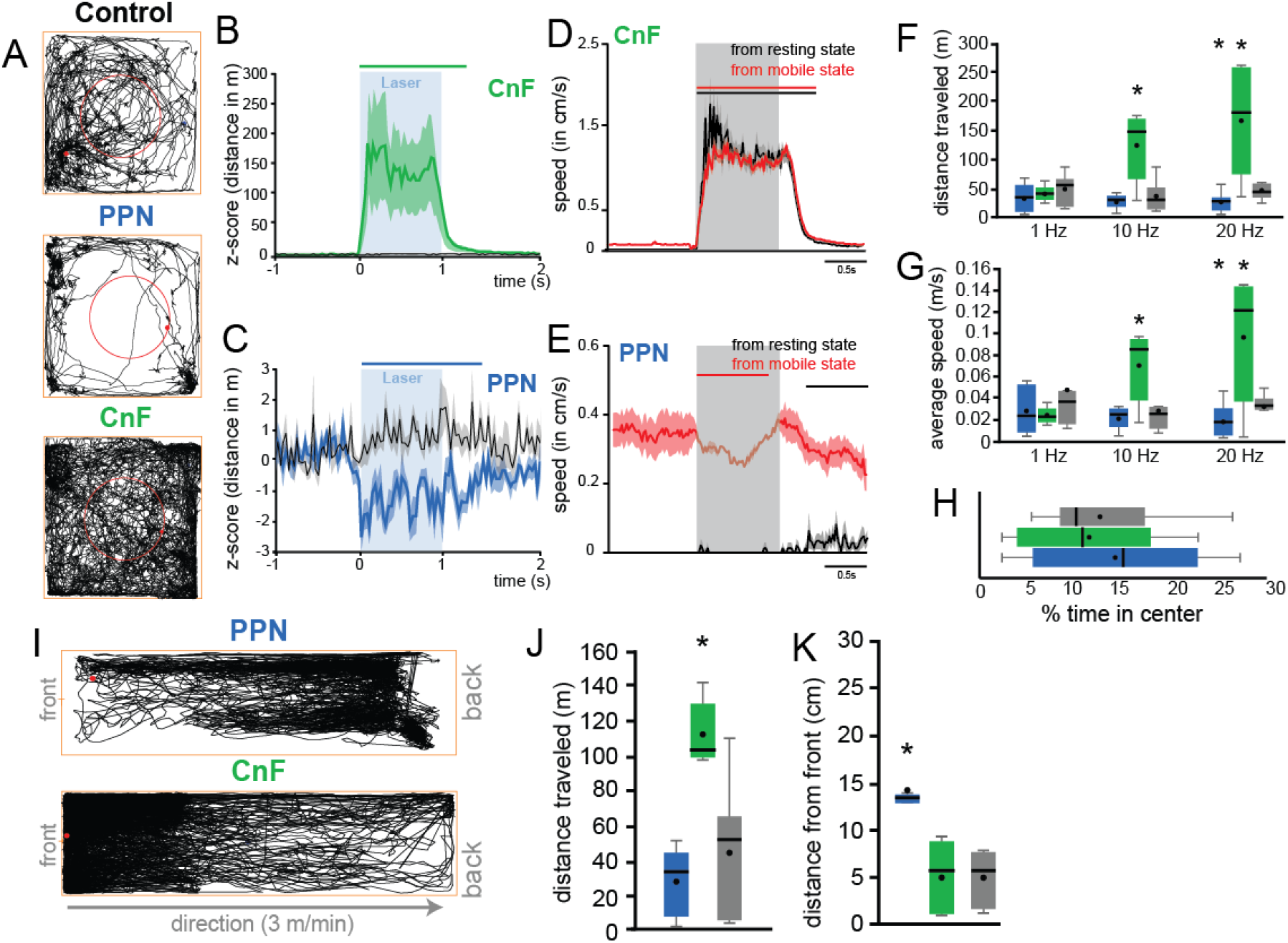
Locomotor effects following stimulation of PPN and CnF glutamatergic neurons. **A,** Trace examples of control, PPN, and CnF stimulated animals tested in the open field. Red circle represents the center of the arena. The behavior was recorded at a resolution of30 frame-per-second. **B-C,** Normalized distance traveled (5ms bin) during individual 10Hz stimulation of CnF (**B**) or PPN (**D**) glutamatergic neurons; control animals (gray; wild-type) received the same experimental treatment (two-way MANOVA groups x stimulation:, group effect F(2,491)=198.21, P=0.00001, stimulation effect F(2,491)=503.43, P=0.00001, interaction effect F(4,491)=201.55, P=0.00001; posthoc Bonferroni, P_CNFstim_PPNstim_=0.0001, P_CNFstim_CTRLstim_=0.0001, P_PPNstim_CTRLstim_=0.0001). The line above represents the statistical difference of the distance traveled compared to the baseline (1s). **D-E,** Distance traveled per 5ms bin during individual stimulation of CnF (**D**) or PPN (**E**) glutamatergic neurons during resting (black) or during spontaneous movement (3-way mixed ANOVA structure x state x stimulation: stim effect F(2,443)=9, P=0.0001; structure: F(1,443)=79.95, P=0.00001, state: F(1,443)=2792.70, P=0.00001, interaction: F(4,443)=30.72, P=0.00001, posthoc Bonferonni P_PPN_mobile_immobile_ = 0.0001; P_cnf_mobile_immobile_> 0.05). The line above represents the statistical difference of the distance traveled compared to the baseline (1s).**F-G,** Total distance traveled and average speed (in m/s) following stimulation at 1Hz, 10Hz or 20Hz (20ms pulse; distance traveled: two-way RM-ANOVA: F_group_(2,62)=0.86, P=0.433, F_frequency_(2,62)=5.22, P=0.0102, F_interaction_(4,62)=6.83, P=0.0003, post hoc: 1Hz: P_ctrl-CNF_=0.256, P_ctrl-PPN_=0.11; 10Hz: P_ctrl-CNF_=0.002, P_ctrl-PPN_=0.25; 20Hz: P_ctrl-CNF_=0.002, P_ctrl-PPN_=0.035; average speed: F_group_(2,62)=2.71, P=0.1046, F_frequency_(2,62)=1.93, P=0.16, F_interaction_(4,62)=5.55, P=0.0014, post hoc: 1Hz: P_ctrl-CNF_=0.11, P_ctrl-PPN_=0.14; 10Hz: P_ctrl-CNF_=0.008, P_ctrl-PPN_=0.25; 20Hz: P_ctrl-CNF_=0.007, P_ctrl-PPN_=0.031). **H**, Percentage of time spent in the center of the arena (F(2,20)=0.30, P=0.7426). **I,** Representative traces of PPN and CnF stimulated animals on the constant-speed treadmill. **J-K,** Distance traveled and average distance from the front of the treadmill following stimulation at 10Hz (distance traveled: one way ANOVA F(2,21)=24.03, P=0.00001, Bonferonni P_PPN_CTRL_=1.0, P_CTRL_CNF_=0.0001, P_PPN_CNF_=0.0001; average speed: F(2,21)=17.41, P=0.0001, Bonferonni P_PPN_CTRL_=0.714, P_CTRL_CNF_=0.0001, P_PPN_CNF_=0.001; distance to the front: (F(2,21)=14.44, P=0.0001, Bonferonni P_PPN_CTRL_=0.0001, P_CTRL_CNF_=0.715, P_PPN_CNF_=0.001). * P<0.05. All experiments have been replicated at least 3 times. Whisker plot are representing mean, median, standard error and 25/75^th^ percentile. All data are represented as mean ± SEM.

To determine whether the reduction in motor activity observed following PPN stimulation was the consequence of altering the behavioral state during exploratory locomotion (i.e. as evaluated above in the open field) or rather a pure motor effect, we next tested the mice during forced locomotion (custom-made motorized treadmill, 3m/min constant speed), in which animals keep up walking at the front of the treadmill (as seen in controls; **Fig. 5I-K**). In the PPN group, blue light stimulation caused the mice to stop locomotion and lag at the rear of the treadmill (**Fig. 5I-K, Supplementary Fig. 5C**). As expected, mice in the CnF group spent most of the time at the front of the treadmill and had a significantly larger traveled distance than control and PPN groups (**Fig. 5I-K**). These results suggest that activation of PPN glutamatergic neurons reduces locomotion by decreasing overall motor activity.

Because the MLR, and specifically the PPN, have been proposed to have a key role in gait and balance, we next tested the mice in the elevated grid walk test, which evaluates skilled coordinated movements requiring sensorimotor integration^43^. Mice were placed on a 20 x 40 cm elevated grid (grid size 1.5 cm) and allowed to explore freely for 20 minutes (**Fig. 6A-B**). Compared to controls, stimulation of PPN neurons produced a significant reduction in the distance traveled, distance to the center of the grid and movement speed (PPN n=6; control n=11; **Fig. 6B-D, Supplementary Fig. 5D**). Furthermore, PPN stimulation increased the number of foot slips (**Fig. 6E-F, Video 2**) whereas the number of rearing events decreased (**Supplementary Fig. 5E**), suggesting a disrupted sensorimotor integration. This effect likely contributed to the markedly reduced exploration of the grid area observed only in mice of the PPN group. In contrast, stimulation of CnF neurons (n=6 mice) resulted in mice jumping off the grid as a consequence of the robust motor activation, despite using lower frequencies and lower laser power, and therefore these experiments were not quantified. Altogether, these results indicate that activation of CnF glutamatergic neurons produced motor responses with no voluntary control, i.e., regardless of the behavioral context, and the intensity of the response was only determined by the frequency of stimulation. On the other hand, activation of PPN glutamatergic neurons blocked distinct components of motor activity regardless of the frequency of stimulation, including locomotion and gait.

**Figure 6.**
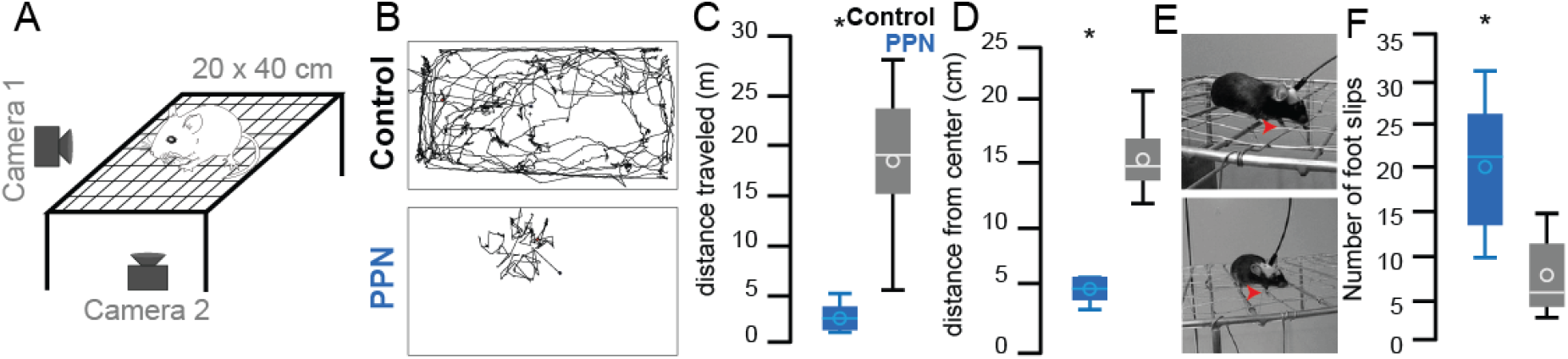
Modulation of gait by PPN glutamatergic neurons. **A,** Representation of the elevated grid walk test. **B,** Representative traces in control and PPN groups (CnF group is not shown, see text for details). **C-D**, Distance traveled (t-test two-tail t(28)=7.8146, P=0.00001) and average distance to the center (t(28)=6.34, P= 0.00001) following 10Hz stimulation. **E**, Representative images of mice making footslips during the elevated grid-walk test. **F**, The total number of footslips (t(15)=5.29, P=0.00001). * P<0.05. All experiments have been replicated at least 3 times. Whisker plot are representing mean, average, standard error and 25/75^th^

### Differential modulation of muscle activity by PPN and CnF neurons

Classically, the function that defines the MLR is the modulation of locomotion. Despite both PPN and CnF providing excitatory innervation to motor structures in the lower brainstem, medulla and spinal cord, the effect of activating each neuronal group separately revealed contrasting effects during motor behavior. To determine whether the seemingly opposing effects on locomotion reflect a competing process between PPN and CnF neurons, or rather a cooperative mechanism to produce an integrated motor output, we measured the impact of each group of neurons on muscles involved in locomotion. VGLUT2-Cre mice were unilaterally transduced with ChR2 in either the PPN (n=4) or the CnF (n=4; control wild-type, n=3) and were bilaterally implanted with bipolar EMG electrodes in both the forelimb and hindlimb biceps. Mice were recorded during spontaneous behavior in their home cage and single blue light pulses (20 ms) were randomly delivered (**Fig. 7A**). First, we measured the effect on the muscle tone and found that blue light stimulation equally induced an increase in the EMG signal in the PPN and CnF groups but followed different dynamics: CnF stimulation transiently increased the RMS signal resulting in a short muscular activation, whereas PPN stimulation produced a long-lasting contraction (response duration in ms: PPN: 1383.59 ± 59.50, CNF: 384.91 ± 15.71, **Fig. 7B-C**). The analysis of the first 500 ms after the onset of the blue laser revealed that the magnitude of the responses between PPN and CnF groups is similar (% of change 0-0.5s, PPN: 193.988+/−13.504, CnF: 213.378+/−29.346, CTRL: −5.24+/−4.21, one-way ANOVA: F(2,214)=4.89, P=0.0084, post hoc Bonferroni P_PPN-CTRL_=0.017, P_PPN-CNF_=0.9, P_CNF-CTRL_=0.006), but the difference becomes evident following this initial phase, denoting a long-lasting effect in the PPN group (% of change 0.5-2s, PPN: 78.60+/−10.06%, CnF: 15.72+/−3.93%, CTRL: 6.66+/−4.55%, one way ANOVA F(2,214)=26.26, P=0.00001, Bonferoni post hoc, P_PPN-CTRL_=0.0001, P_PPN-CNF_=0.0001, P_CNF-CTRL_=0.04, **Fig. 7B-C**). Furthermore, the response latency following PPN stimulation was significantly shorter than in the CnF group (PPN: 28 ± 4.78 ms, CNF: 79.45 ± 10.58 ms; two-tailed t-test t(211)=−3.58, P=0.0004; **Fig. 7D)**. These results suggest that PPN stimulation produces a robust and long-lasting effect on the muscle tone, contrasting with a short-lasting effect that follows the stimulation of the CnF.

**Figure 7.**
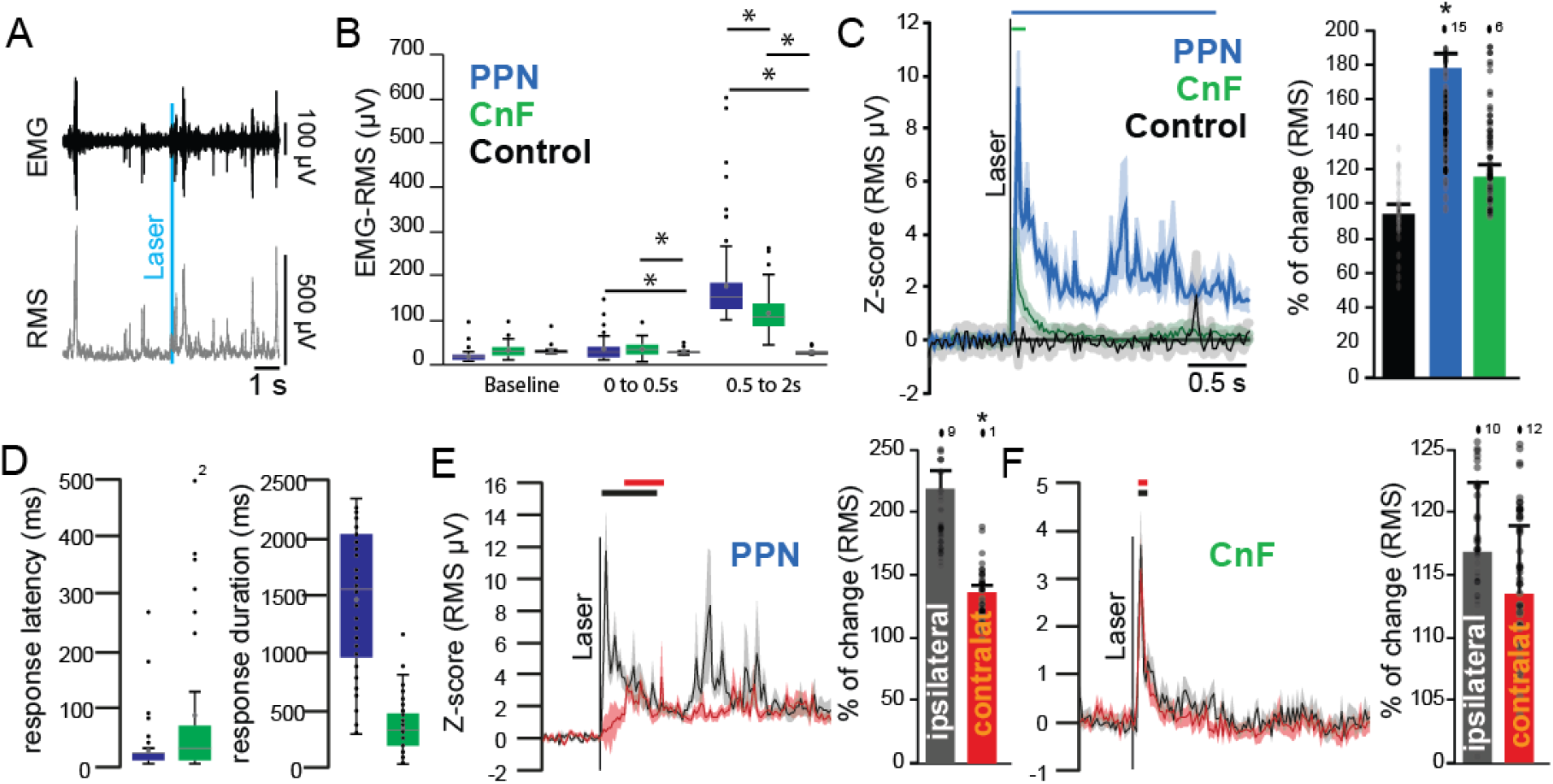
Differential involvement of PPN and CnF glutamatergic neurons in muscle tone generation. **A,** Example of electromyogram (EMG) activity recorded at the level of the biceps following laser stimulation and the conversion into root-mean-square (RMS). **B**, Raw amplitude (μV) of the RMS-EMG of ipsilateral forelimb biceps during baseline, immediately or 500 ms after single stimulation pulses delivered in the PPN, CnF or sham (0-0.5s after stimulation: one-way ANOVA F(2,214)=5.59, P = 0.0001, post hoc Bonferroni P_PPN_vs_CTRL_=0.004, P_CNF_vs_CTRL_=0.004, P_PPN_vs_CNF_=1.0; 0.5 to 2s after stimulation: F(2,214)=46.62, P=0.00001, post hoc Bonferroni P_PPN_vs_CTRL_=0.0001, P_CNF_vs_CTRL_=0.0001, P_PPN_vs_CNF_=0.001). **C**, Change in the RMS signal following repeated single stimulation pulses recorded at the level of the biceps (% change relative to 1s baseline: PPN: 178.60±10.06%, CNF: 115.72±3.93%, CTRL: 94.33±4.55%, one-Way ANOVA F(2,214)=26.26, P=0.00001; Post hoc Bonferroni P_PPN-CNF_=0.0001, P_PPN-CTRL_=0.0001, P_CNF-CTRL_=0.804). **D**, Response latency and duration of the significant increase in the RMS signal in response to PPN or CnF stimulation (latency: F(1,212)=6.44, P = 0.019; duration: F(1,212)=19.29, P=0.00001). **E-F**, Change in the RMS signal in the ipsilateral and contralateral forelimbs biceps following stimulation in PPN (PPN_ispi_=218.08±17.02%, PPN_contra_ 138.46±5.71%, Two-Way ANOVA stim x side: F_stim_(1,401)=648.221, P=0.00001, F_side_(1,401)=39.6, P=0.0001, F_interaction_=242.6, P=0.00001) and CnF groups (CnF_ispi_: 117.06±5.54%, CnF_contra_: 113.55±5.51%, two-Way ANOVA stim x side: F_stim_(1,401)=51.29, P=0.00001, F_side_(1,401)=1.80, P=0.18, F_interaction_(3,401)=18.30, P=0.0000). Lines represent statistical difference compared to baseline. * P<0.05. All experiments have been replicated at least 3 times. Whisker plot are representing mean, median, standard error and 25/75^th^ percentile; individual data are represented by small dots. Out of range data points are reported as numbers above the histogram. All data are represented as mean ± SEM.

We next evaluated the effect of the stimulation on the contralateral musculature. Stimulation of PPN neurons produced a marked increase in the amplitude of the ipsilateral biceps that was significantly larger than the contralateral biceps. In contrast, stimulation of CnF neurons produced similar increases in the EMG amplitude of the ipsilateral and contralateral biceps (**Fig. 7E-F, Supplementary Fig. 6**). Thus, while activation of PPN neurons produces a long-lasting increase in the amplitude of the ipsilateral EMG consistent with an increased muscle resistance to passive movement, the bilateral nature of the short-lasting muscle activation observed after unilateral CnF stimulation is consistent with the frequency-dependent bouts of locomotor activity. Further evidence was obtained following CnF low- (0.1Hz) or high-frequency (5Hz) stimulation while mice were held in a tail-lifted position revealing motor contractions that resemble context-independent involuntary locomotion (**Video 3**). In contrast, the lasting increase in muscle tone observed following PPN stimulation may act as a readiness signal that precedes locomotion, suggesting that both MLR structures act in coordination to modulate the motor output. These results, together with the differences in connectivity and physiological properties, uncover fundamental differences in the modulation of muscle activity by MLR neurons and reveal their differential roles in motor behavior.

## Discussion

Neurons of the MLR have been classically identified as a critical node for the integration of behavioral signals originating in forebrain systems related to the modulation of motor output. The results presented in this study reveal several differences between MLR substructures in terms of their connectivity, physiological properties and effects on motor behavior and muscle activity. In terms of connectivity, we show that PPN neurons have widespread projections to a variety of motor regions including the basal ganglia and spinal cord, whereas CnF neurons mainly concentrate in the brainstem. In terms of physiological properties, we show that PPN neurons comprise a heterogeneous group displaying a range of adapting responses, whereas the majority of CnF neurons are fast-adapting. In terms of behavior, we show that stimulation of PPN neurons decreases overall motor activity whereas CnF stimulation produces robust and highly-reliable bouts of motor activity. Finally, stimulation of PPN neurons produces a prolonged increase in muscle tone whereas stimulation of CnF neurons produces brief, bilateral motor contractions of the limbs. Thus, the distinct attributes observed among MLR structures reveal major differences in their composition and properties, and shed light into the fundamental mechanisms underlying their role in motor behavior.

Our data reveal that CnF glutamatergic neurons control a stereotypical motor response that scales its intensity with optogenetic frequency; from high-velocity locomotion to jumping, CnF stimulation causes rapid movement of the hindlegs independent of context. Subsequently, we observed that these motor effects can be explained by fast, monophasic and bilateral muscle responses that mostly occurred within the 0.25 seconds immediately following optogenetic CnF stimulation, in contrast with PPN stimulation which caused multiphasic EMG fluctuations above baseline for an average of 1.5 seconds, sometimes longer than 2 seconds. Along with the swift and consistent effect of CnF stimulation on muscle and motor responses, we found that the majority of CnF neurons (85.7%) are fast-adapting and strongly accommodating, suggesting that they are capable of generating phasic motor responses in response to the synaptic drive by upstream structures. Predominant inputs to the CnF are the superior colliculus, inferior colliculus, and the periaqueductal gray area, providing a basis for the rapid transmission of sensory information in contexts that signal threat. Along these lines, the only regions that showed a higher density of synaptophysin-positive axons from the CnF than the PPN were the hypothalamus (notably, the preoptic nucleus) and the habenula, regions involved in homeostatic regulation^44,45^ and the valuation of threat^46,47^, respectively. Overall, our data support a developing theory that the CnF is involved in fast-escape behavior^7,48^ and its activity is likely to be modulated to fast-incoming sensory information.

In contrast to the CnF, we show that PPN glutamatergic neurons display heterogeneous features as revealed by a wider input/output connectivity map, a range of spike adaptation profiles, and distinct effects on motor behavior. Whereas the CnF exhibited a more restricted output domain, synaptophysin-positive PPN axons were observed in the spinal cord, medulla, midbrain, cerebellum, thalamus, basal ganglia and cortex. Notably, every single brain region that provides input neurons projecting to the CnF also provides input to the PPN. Of the neurons we recorded in the PPN, 48.8% were fast-adapting neurons, 30.2% non-adapting neurons, and 21% slow adapting, suggesting a greater diversity of neuronal profiles than the CnF. In terms of behavior, PPN stimulation causes stopping in the open field and on the treadmill, with no significant relationship to the 10Hz and 20Hz stimulation frequencies used. These findings agree with prior studies that have shown decreased locomotion^12^ or no increase in locomotion^7^ due to PPN stimulation at these frequencies (but see^7^ for the effect at higher frequencies). Furthermore, we found that stopping behavior was not the only PPN-dependent phenomenon observed. For instance, on the elevated grid walk test, PPN stimulation led not only to decreased travel distance and more time in the center of the grid, but also significantly more foot slips, which could be interpreted in the context of a loss of motor coordination (as seen in lesions to PPN cholinergic neurons^49^) or reset of the motor action sequence (potentially through activation of striatal interneurons^33^). Nevertheless, an alternative mechanistic interpretation based on the EMG data suggests that PPN neurons increase the muscle tone in preparation for movement, but they lack the capability of triggering a motor output by themselves. Compared to CnF-derived muscle responses, PPN stimulation caused EMG activations that were multiphasic and 6-to-8 times longer on average, and produced increases from baseline that were 3 times more significant. Notably, while the short-lived CnF muscle activation was tightly correlated with locomotion bouts, PPN muscle activation was not. Thus, the effect of PPN stimulation on the EMG revealed a prolonged increase in muscle tone in the absence of movement that is consistent with the muscle preparation that would be necessary to execute upstream-driven (e.g. basal ganglia) motor commands. Such interpretation is congruent with the activity of PPN glutamatergic neurons in arousal and behavioral activation^25,50–52^, suggesting that PPN neurons encode a readiness signal that enables motor responses. Altogether, our results uncover new aspects of the heterogeneity observed in PPN glutamatergic neurons^24^ which will most certainly hold important clues to understand their multifarious contributions to behavior, and highlight the necessity of future studies to address this in detail.

Growing evidence show that the function of PPN neurons is closely linked to the basal ganglia. For example, PPN glutamatergic neurons are capable of reliably patterning dopamine release via synapses targeting the soma, proximal dendrites, and axon initial segment of SNc dopamine neurons^53^. Furthermore, PPN glutamatergic neurons innervate striatal interneurons and produce feed-forward inhibition of the striatal output^33^. In addition to the SNc and the striatum, our data revealed synaptophysin-positive axons in the globus pallidus, endopenduncular nucleus, SNr and VTA originating in the PPN. In comparison, the CnF only projects to the VTA, SNr, and globus pallidus. Altogether, basal ganglia structures receive a greater density of axons from the PPN than the CnF. The PPN also exclusively targets the CM-Pf thalamus, which exhibit strong control on the basal ganglia by gating input to the striatum prior to the selection of goal-directed actions^54,55^, signaling saliency^56^, and controlling the learning of new action-outcome contingencies^57^ by affecting striatal microcircuitry via control of specific interneuron subpopulations^57,58^. In terms of its afferent connectivity, our data shows that every single node of the basal ganglia provides an input to the MLR (predominantly to the PPN). In particular, neurons of the SNr, which constitute the main basal ganglia output in rodents, is one of the primary structures providing inputs to PPN (but not CnF), as revealed by ourselves and others^6,7^. Nevertheless, despite the close bidirectional connectivity and functional analogy between PPN and the basal ganglia, our data reveal that the input/output connectivity map of PPN glutamatergic neurons is far more distributed and intricate than previously considered. This suggests that a number of other brain regions may converge on basal ganglia output-recipient PPN neurons, thus conferring them with the potential to weigh the distinct synaptic inputs and select an integrated behavioral output.

In the past decade, the PPN has emerged as a potential target for deep brain stimulation (DBS) with a mixture of results thus rendering its use controversial^55–57^. Our work suggests that the variability observed in the clinical setting may partly be due to differences in electrode location and/or stimulation frequency and intensity. Although investigations of the two excitatory structures comprising the MLR provide insight toward a general model, the complexity of the MLR input/output map suggests a topography of domain-specific subnetworks that must be examined specifically to interpret the variability observed following PPN-DBS in clinical populations. The variety of observed motor effects due to MLR stimulation between our study and others^6,7^ is likely a manifestation of different PPN sub-circuits being recruited due to varying experimental manipulations (i.e. fiber optic location, extension of the ChR2 transduction area and/or stimulation frequency and intensity). One possibility could be that the strong effects that the PPN has on dopamine release are only recruited under specific stimulation parameters and provide a basis for exploratory locomotion (e.g.^59^) whereas other PPN glutamatergic circuits modulate muscle tone. An alternative explanation is that over-recruitment of segregate PPN pathways by optogenetics results in stopping. Pathway-specific interventions controlled by stimulation site, frequency, or intensity could provide a new dimension by which to analyze the MLR as a versatile DBS target.

## Methods

### Animals

Homozygous floxed-tdTomato (B6;129S6-Gt(ROSA)26Sortm9(CAG-tdTomato)Hze/J; Jax number: 007905), VGLUT2-cre (Slc17a6tm2(cre)Lowl(also called VGLUT2-ires-Cre); Jax number: 028863), and wild-type (C57bl/6, Jax number: 000664) adult male and female mice were used for all experiments. All mice were housed on a normal 12:12h light:dark cycle (light on at 7:00) and had unrestricted access to food and water. All experiments were performed in accordance with the National Institutes of Health Guide to the Care and Use of Laboratory Animals, or the Hungarian and International EU Directive 2010/63/EU for all animal experiments. Approval was obtained from Rutgers University Institutional Animal Care and Use Committee (16054A1D0819) and the Committee of Animal Research of the University of Debrecen (5/2015/DEMAB).

### Viral Injections/Surgery

All surgeries were performed under aseptic conditions. Body temperature was maintained at 37±1 degree C using a heating pad. Mice were deeply anesthetized with isoflurane (1.5% to 4%, in O_2_) and placed in a stereotaxic apparatus (David Kopf Instruments). Ophthalmic ointment was applied. Following skin incision, a small cranial hole was made above the targeted area. All measurements were made relative to bregma and dorsoventral coordinates were set from dura. Viral injections were performed using a 32-gauge syringe (Hamilton Syringes Neuros, #65458) at 5-7 nl/min rate using a microsyringe pump (micro4, WPI). For multiple injections, an additional syringe was used to avoid contamination, and syringes were thoroughly cleaned with ethanol and water between each experiment. After completion of the injections, 10 to 15 min were allowed before slowly withdrawing the syringe. At the end of the surgeries, animals received injections of Buprenorphine (0.10mg/kg, sc) and Baytril (0.05mg/kg). Viruses used for all experiments were as follows: AAV2-DIO-EF1*α*-YFP (titer: 10^12; injection in PPN: 20nL; injection in CnF: 15nL, UNC Vector Core); AAV2-DIO-EF1*α*-YFP-2A-synaptophysin-mRuby (titer: 10^12; injection in PPN: 10nL; injection in CnF: 10nL, Stanford Vector Core); AAV5-DIO-TVA-mCherry (titer: 10^12; injection in PPN: 10nL, injection in CnF: 7.5nL, UNC Vector Core); AAV8-DIO-RG (titer: 10^12; injection in PPN: 10nL, injection in CnF: 7.5nL, UNC Vector Core); RvDG-YFP (titer: 10^8; injection in PPN: 200nL, injection in CnF: 200nL, Salk Institute), AAV2-Flex-EF1a-ChR2(H134R) (titer: 10^12; injection in PPN: 20nL, injection in CnF: 15nL UNC Vector Core). Injection were delivered in the following coordinates (in mm from Bregma): PPN: AP: −4.5, ML: ±1.25, DV: 3.3; CnF: AP: −5.0, ML: ±1.2, DV: 2.2.

### Histology

After *in vivo* experimental procedures were completed, animals were deeply anesthetized with sodium pentobarbital (200 mg/kg) and transcardially perfused with 20ml of phosphate buffer solution (PBS 0.05M) followed by 20ml of paraformaldehyde (PFA 4%). The entire brain and the spinal cord were removed and post-fixed in PFA for 12h. Before slicing, the brain was embedded in a single block of Agar (in PBS, 2%) as well as 3-5mm sections of the spinal cord that were collected in anteroposterior order. Spinal cord and brain sections were sliced at 50μm following coronal or sagittal axes and collected in individual well-plates with 300μm spacing between consecutive sections. All immunohistochemistry solutions were prepared in a solution of PBS with 0.3% Triton (PBS-Triton). First, sections were blocked in PBS-Triton containing 10% normal donkey serum (NDS, Jackson Immunoresearch) for 1h at room temperature, following 3-5 washes with PBS, sections were transferred in a primary antibody solution containing PBS-Triton, 1% NDS and the corresponding primary antibody. The primary solution was left overnight at 4h under constant gentle shaking. Sections were then washed 3-5 times with PBS before to be transferred to the secondary antibody solution (PBS Triton, 1% NDS, and the corresponding secondary antibody) and kept under constant gentle shaking for 4-5h at room temperature. Sections were then washed 3-5 times in PBS before mounted on microscope slides using a mounting medium (Vectashield) and prepared for imaging. Primary antibodies were as follows: mCherry (used for mRuby, mCherry and TdTomato, made in mouse, monoclonal, ABCAM AB167477, concentration 1:1000), ChAT (choline acetyltransferase, made in goat, polyclonal, Merk Millipore, AB144P, concentration 1:500), GFP (to enhance eYFP detection, made in rabbit, polyclonal already conjugated-488, Thermofisher, A21311, concentration 1:1000) and Fluorogold (made in rabbit, polyclonal, Merck Millipore, AB153-I, concentration 1:1000). Secondary antibodies were as follows: anti-Goat CY3 (raised in donkey, Jackson Immuno-research 705-165-147, concentration 1:1000); anti-Goat CY5 (raised in donkey, Jackson Immuno-research 705-175-147, concentration 1:1000), anti-mouse CY3 (raised in donkey, Jackson Immuno-research 715-165-150, concentration 1:1000), anti-rabbit 488 (raised in donkey, Jackson Immuno-research 711-545-152, concentration 1:1000) and anti-rabbit AMCA (raised in donkey, Jackson Immuno-research 711-155-152, concentration 1:1000).

### Imaging

Fluorescent images were captured using a confocal laser microscope (Olympus FV1000S) with the FluoView software (Olympus), under a dry 10X/0.40 NA objective, 20X/0.40NA or an oil-immersion 63X/1.40NA objective. All sections were first acquired at high resolution (10X, 1024 * 1024 pixels) using mosaic reconstruction to determine the virus diffusion, the viral injection site and the placement of the optic fiber. For cell counting, sections were scanned at 20X using medium resolution (1048 * 720). For projections and synapses counting, sections were acquired at high resolution (20X, 2048 *2048 pixels), with a 1μm-optical section z-stack across 40μm (top and bottom 5μm of the section were discarded). Single images of axonal projections or synaptic contacts were acquired at high magnification (63X), high-resolution (2048*2048 pixels), with 4-time deconvolution and a 1μm-optical section z-stack across 40μm. All pictures were saved as images and metadata in order to correct the mosaic alignment using Photoshop (version 5, Adobe). All fluorescent images were transferred to Fiji software, were color-converted based on the secondary antibody and the filter used (AMCA: 400 – 450 nm, Alexa488: 500-550 nm, CY3: 590-620 and CY5: 650-700), signal-adjusted, and merged using in-built tools.

### Cell counting

Each brain and spinal cord section scanned were converted into bitmap images, duplicated and overlapped with the outline of the Paxinos and Franklin^60^, mouse brain Atlas (7th edition). Images were then transferred to Fiji, and in-built counting tools were used. The number of cell markers per nucleus (as defined by the Atlas) was then transferred to an Excel spreadsheet. The counted cells of each identified brain structure that were represented in separate sections were put together for the final analysis and normalized to the total number of neurons counted in all brain sections collected for each animal. For RvdG experiments, the YFP-positive neurons located in the site of injection (PPN/CnF) were not quantified for the whole brain mapping. For the spinal cord, a random number of spinal cord sections that were representative of all segments (40-80 sections per mice) were processed and counted. Based on the average number of inputs neurons per section found, we extrapolated the putative number of inputs in the entire spinal cord using an average length of 3.2cm. Normalized data and raw data were tested for normality and compared using a Wilcoxon rank-sum test (non-parametric). The threshold to significance was determined at P<0.05. All data were shown as mean±SEM.

### Synapse density estimation

Brain sections were prepared as above. The nucleus’ outlines were drawn using built-in tools and the number of pixels above the threshold, the surface area, the background gray value, and the average gray value within the drawing area were obtained to define the density of the synapses using the formula:

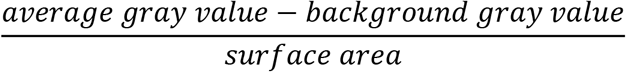

The area that was considered as an artifact due to dust or air-bubbles generated during immunohistochemistry or slice mounting was manually discarded using a similar approach. Normalized data and raw data were tested for normality and compared using a Wilcoxon rank-sum test (non-parametric). The threshold to significance was placed at P<0.05. All data were shown as mean±SEM.

### Ex Vivo Electrophysiology

9-16 days old animals expressing tdTomato fluorescent protein in a VGLUT2-dependent way (n = 25) were used for the slice electrophysiology experiments. Coronal midbrain slices (with 200 μm thickness) were prepared in low Na+ aCSF (cca. 0 - −2 °C) with a Microm HM 650V vibratome (Microm International GmbH, Walldorf, Germany). The slices were incubated in normal aCSF for 1 hour on 37°C prior to starting the experiment. The resistance of the patch pipettes was 5-7 MΩ, and the composition of the internal solution was the following (in mM): K-gluconate, 120; NaCl, 5; 4-(2-hydroxyethyl)-1-piperazineethanesulfonic acid (HEPES), 10; Na2-phosphocreatinine, 10; EGTA, 2; CaCl2, 0.1; Mg-ATP, 5; Na3-GTP, 0.3; biocytin, 8; pH 7.3. Whole-cell patch-clamp experiments were conducted at room temperature with an Axopatch 200A amplifier (Molecular Devices, Union City, CA, USA). Clampex 10.0 software (Molecular Devices, Union City, CA, USA) was used for data acquisition, while data analysis was performed by Clampfit 10.0 (Molecular Devices) software. Only stable recordings with minimal leak currents were considered, and only recordings with series resistance below 30 MΩ, with less than10% change, were included. Both voltage- and current clamp configurations were employed. Protocols and recorded parameters are represented in Supplementary Table 1. In certain experiments, 1 μM tetrodotoxin (TTX; Alomone Laboratories, Jerusalem, Israel) was administered to eliminate action potential generation in the preparation. Visualization of the genetically encoded fluorescent marker (tdTomato) was achieved by using a fluorescent imaging system (Till Photonics GmbH, Gräfeling, Germany) containing a xenon bulb-based Polychrome V light source, a CCD camera (SensiCam, PCO AG, Kelheim, Germany), an imaging control unit (ICU), and the Till Vision software (version 4.0.1.3).

#### Morphological analysis of the recorded neurons

Patched neurons were labeled with biocytin and samples were fixed (4% paraformaldehyde in 0.1 M phosphate buffer; pH 7.4; 4 °C) for morphological identification of the neurons. Tris-buffered saline (in mM, Tris base, 8; Trisma HCl, 42; NaCl, 150; pH 7.4) supplemented with 0.1% Triton X-100 and 10% bovine serum (60 min) was used for permeabilization. Incubation was performed in phosphate buffer containing streptavidin-conjugated Alexa488 (1:300; Molecular Probes Inc., Eugene, OR, USA) for 90 min. The cells were visualized using a Zeiss LSM 510 confocal microscope (Carl Zeiss AG). The reconstruction of neurons was performed by NeuroLucida software (MBF Bioscience, Williston, VT, USA).

### Behavioral Assays

#### Laser stimulation

A blue laser (450nm, OEM Laser system) was used to excite ChR2. Stimulation parameter varied. Several stimulation frequencies were used (1Hz-20z) and controlled by a low-noise shutter (SH1, Thorlabs) plugged to the control cube (KSC101, Thorlabs) which in turn was triggered by TTL signals delivered by the Anymaze interface. The laser output was set to be 2-3 mW at the end of the patchcord.

#### Small Open field

Following implantation of the optic fiber, animals were allowed to recover for 5 to 7 days. Animals were then habituated for 5 minutes to the open field before testing. The custom-made open field was developed as following: a dark cube of 40 × 40 × 40 cm with the floor covered with a non-reflective white surface to allow better contrast between the background and the mouse, 4 white lamps were positioned on the top of the cage to allow optimal illumination. Animals were tested for 30 minutes, with optogenetic stimulation 1s ON/9s OFF (20ms pulses, <3mW laser power) using stimulation frequencies of 1, 10 or 20Hz. The animal movements were recorded using a high-speed/high-resolution camera (120fps) RunCam2 and the software Anymaze (Stoelting). The software was tracking the gravity center (body) of the animal, the head and the tail position. The time in the center (15cm circle located in the center of the field) or periphery was defined based on the position of the animal body. The software recorded the animals’ speed, distance traveled, time in center and time in the periphery. Whilst online analyses of the above-mentioned parameters were based on 30 recorded frames per second, offline analyses used 120 frames per second. On- and offline analyses were compared and sequences differing in more than 5% were discarded. Stimulation delivery was controlled using the software interface Ami1.

#### Large open field

The above experiment was repeated in a larger open field, which consisted of a dark cube of 80 × 80 × 80 cm with the same features as the one described above. A small slope was built at the base of each wall to avoid animals making contact with the walls. Animals were tested for 60 minutes, optogenetic stimulations were delivered for 1 minute (repeated loops of 1s ON/9s OFF on) and were spaced by 1 minute with no stimulation. Stimulation protocol was as follow: 20ms pulses, <3mW laser power and the frequency was increased from 0.1 to 30 Hz (0.1, 0.5, 1, 2.5, 5, 7.5, 10, 12.5, 15, 17.5, 20, 22.5, 25, 27.5, 30). Animals movement were recorded and analyzed as above.

#### Treadmill

The treadmill apparatus consists of a custom-made belt of 10cm by 30cm that is operated at constant speed. The animals’ position, the center of the body, head, and tail were monitored using the Anymaze software via a high-speed camera (RunCam2) camera placed above the treadmill. Further, the behavior was also monitored using a camera located on the side of the treadmill. Animals were tested for 15 minutes while receiving optogenetic stimulation (1s ON/9s OFF, 20ms pulses, <3mW, 10Hz). 2 days before testing, animals were habituated to the still treadmill for 10 minutes. One day before testing, the habituation occurred on the moving treadmill. On the day of testing, animals were connected to the laser and placed on the treadmill with the speed set at 3m per minute. Animals were tracked and analyzed to determine distance traveled, average speed and position of the head-body and tail-body axes. Stimulation delivery was controlled using the software interface Ami1.

#### Elevated grid

The elevated grid apparatus consists of a custom made 60 × 30 cm grid (1.5 × 1.5 cm grid space), elevated 1m off the floor and illuminated from the bottom. The animals’ behavior was monitored by two high speed-cameras located on the side (for rearing and foot slips) and on the bottom (for the animal position). Animals were not exposed to the apparatus before testing to avoid any habituation, but all animals were handled for several days before testing. On the day of the testing, animals were connected to the laser, placed in the middle of the grid and their behavior was monitored for 20min using Anymaze software while receiving optogenetic stimulation for 1s every 9s (20ms pulses, <3mW, 10Hz). Animals were tracked and analyzed to determine distance traveled, average speed and position of the head-body and tail-body axes. Stimulation delivery was controlled using the software interface Ami1.

#### Control animals

For each experiment, control animals consisted of WT animals receiving the same manipulations and undergoing the same procedures as the experimental groups. Control animals were excluded if the injection was “out-of-target” or the implantation was not correctly positioned.

#### High-resolution analyses

The cartesian coordinates of each acquired frame (120 fps) were converted offline into interframe distance traveled. The peristimulation distance traveled was defined as the “stimulation locked distance traveled” using z-score transformation normalized into the 5s baseline prior to each stimulation. Due to camera fps variability, the cartesian coordinates were used at 5ms intervals by extrapolating the interframe position.

#### Data analyses

All experiments were randomly organized, and data of each animal were analyzed similarly. To prevent data loss during animal tracking, the data from online and offline were compared, and the portion of data was removed if we found any differences in the recording. Following comparison of the online/offline tracking, data were expressed as the following parameters: overall distance traveled, average speed during the entire session, number of ipsilateral of contralateral rotation and distance traveled 5 seconds before and after each stimulation (with 5ms bin size). High-resolution data were converted to z-score of the distance traveled compared to the baseline (−5 to 0s before stimulation). All data were compared between groups using one-way ANOVA or by comparing the frequency of stimulation and groups using multivariate ANOVA. A significant ANOVA effect was compared using Bonferroni posthoc analyses.

### Electromyogram recordings

During a surgical procedure as described above, an incision was made at the neck, forelimbs and hindlimbs of the animals and muscles were exposed. EMG bipolar electrodes were implanted in the biceps brachia and biceps femoris of the ipsilateral and contralateral limbs and the connector was affixed on the skull of the animals. In addition, an optic fiber (flat-cut, 200μm, 0.50NA) was implanted above the PPN or CnF (300μm above the injection site) following viral injections and maintained in position using anchor screws. EMG signals were converted into RMS signal and each trial was analyzed individually. All animals received same number of stimulation to avoid overrepresentation.

### Histological verification

Following staining of sections located on in the vicinity of the injection sites for GFP and ChAT, high-resolution images were acquired and processed using Fiji. All ChAT-positive neurons located at the border of the PPN were labeled using in-built tools, then all YFP-positive cell bodies were labeled and their location recorded. The number of YFP-positive neurons located further than 100μm from the closest ChAT-positive neurons (for PPN and the ventral border of the CnF), or within the colliculus (for the dorsal border of the CnF) was calculated as a percentage of the total number of neurons within the injection site. If more than 5% of YFP-positive neurons were located further than 100μm (for PPN) or closest to 100μm or inside the colliculus (for CnF) the animal was excluded from further analyses.

### Statistical Analyses

Anatomical, in vitro and in vivo data (including behavioral data) are represented as mean±SEM. No power analyses were conducted prior to the experiments and group sizes were determined following comparable previously published experiments. Anatomical data was compared using the Wilcoxon rank-sum test following prior determination of the violation of the assumption of normality of the data. In vitro data was analyzed using Student’s t-test, one-way ANOVA or mixed ANOVA. One way ANOVAs and MANOVAs were conducted for in vivo and behavioral experiments. All ANOVAs were followed by Bonferroni corrected post-hoc tests. Level of significance was set at p<0.05.

## Acknowledgments

This research was supported by an NIH grant NS100824 (J.M.S.), a NJ-DOH grant CSCR20IRG008 (J.M.S.), a NARSAD Young Investigator Award (J.M.S.), the Hungarian National Brain Research Program (B.P.), the OTKA Bridging Fund of the University of Debrecen (B.P.) and Rutgers University. The authors are grateful to Dr. Péter Szücs (Department of Anatomy, Histology and Embryology, Medical Faculty, University of Debrecen) for providing access and for his kind help in Neurolucida reconstructions, and Prof. Miklós Antal (Department of Anatomy, Histology and Embryology, Medical Faculty, University of Debrecen) for providing VGLUT2-cre mice for breeding. We also thank Dr. Nadine Gut for comments on this manuscript.

## Data Availability

All custom script, unprocessed figures, whole-brain scans, recordings data are available under reasonable requests.

## Supplementary material

**Supplementary Figure 1.Histological analysis. Related to Figures 2, 3, 5 and 6.**

**A,** Virus spread (circles) and locations of the tip of the optic fibers (red circle) for PPN (green) and CnF (blue) groups. **B,** Fluorescent micrographs of PPN glutamatergic neurons expressing GFP and synaptophysin. **C,** High-resolution images of a transduced CnF glutamatergic axon in PAG expressing GFP in the shaft and boutons and synaptophysin-mRuby in the terminals.

**Supplementary Figure 2. Input/output relationship of PPN and CnF. Related to Figures 2 and 3.**

**A**, Schematic summary of PPN (blue) and CnF glutamatergic (green) axonal distribution using relative synaptic density. **B**, Schematic summary of PPN (blue) and CnF glutamatergic (green) neuron inputs. **C-D,** Graphical representation of inputs and outputs of PPN and CnF glutamatergic neurons based on the data presented in Figures 2 and 3. Input arrows (left) are defined based on the normalized distribution of inputs neurons. Output arrows (right) are defined based on the normalized distribution of the synapses.

**Supplementary Figure 3. Membrane properties of the PPN and CnF glutamatergic neurons. Related to Figure 4.**

**A,** A-current was observed on most PPN and CnF glutamatergic neurons. Current traces elicited by +20 mV voltage step, preceded by −120 mV (black) and −10 mV (red) voltage steps (example shows PPN). The left current trace is the difference of the black and red current traces. **B-D,** Representative examples of the firing properties of PPN glutamatergic neurons. Trains of action potentials elicited by 100 pA depolarizing current injection from −59 mV (**B**), −87 mV (**C**) and −73 mV resting membrane potential (**D**, in the presence of TTX; the arrow indicates the lack of delay). **E-I,** Depolarization and action potential firing elicited by 30 pA depolarizing square current injection from −66 mV (**E**) and −83 mV (**F**) resting membrane potentials. Note the low threshold depolarizing spike (black arrow). (**G**) 30 pA hyperpolarizing current injection from −53 mV resting membrane potential revealed rebound spike and firing (black arrow). **H-I,** Low threshold spike and rebound depolarizing spike (respectively; arrow) in the presence of TTX. **J,** Distributions of functional neuronal types in the PPN and CnF. Group I neurons display low threshold depolarizing spikes but lack A-current (PPN: 22.7%, CnF 33.3%). Group II neurons display A-current (PPN: 47.7%, CnF 37.5%). Group III neurons display both (PPN: 9.1%, CnF 16.6%). Group IIIK lacks all (PPN: 20.5%, CnF 12.5%). **K,** Proportion of PPN and CnF neurons displaying A-current. **L,** Proportion of PPN and CnF neurons displaying low threshold spikes (LTS). All experiments have been replicated at least 3 times. All data are represented as mean ± SEM.

**Supplementary Figure 4. Physiological properties of MLR glutamatergic neurons. Related to Figure 4.**

Statistical summary of the number of action potentials elicited by 1-s depolarizing step, the adaptation index, the ratio of the amplitude of the last and first action potentials of the train, the ratio of the width of the last and first action potentials of the train, the frequency and the duration of the train in different functional subgroups (non-adapting, red; slowly adapting, purple; rapidly adapting, black). The significance was calculated between the first and last datapoints within each trace. * P<0.05, **P<0.01, ***P<0.001. All experiments have been replicated at least 3 times. All data are represented as mean ± SEM.

**Supplementary Figure 5. Frequency-dependent modulation of locomotion in the CnF. Related to Figures 5 and 6.**

**A-B,** Distance traveled following optogenetic stimulation (1s ON/9s OFF) during and immediately after optogenetic stimulation of CnF glutamatergic neurons using a randomized stimulation protocol ranging from 0.1Hz to 30 Hz (mixed ANOVA; *during*: F(14,89)=2.69, P=0.003, trendline: R^2^ = 0.4774, Max: 12.5, Posthoc Bonferroni P=0.034; *after*: F(14,89)=1.24, P=0.2994). Gray dots represent individual data points, black dots represents average values, vertical lines represent SEM, and the red line represents the best fitted trendline (y=−0.0314x^2^+0.7389x+4.9582). **C,** Average speed of animals in the CnF (blue), PPN (green) and control groups (gray) during the treadmill test (F(2,21)=17.41, P=0.0001, Bonferonni posthoc P_PPN_CTRL_=0.714, P_CTRL_CNF_=0.0001, P_PPN_CNF_=0.001). **D,** Average speed of mice in the PPN (green) and control groups (gray) during the elevated grid walk test (t-test two tail: t(28)=6.39, P=0.00001,). **E**, Total number of rearing events observed in the elevated grid walk test following stimulation of PPN glutamatergic neurons (green) and compared to control mice (gray; t(15)=2.63, P=0.0095).

**Supplementary Figure 6. Complementary EMG experiments. Related to Figure 7.**

**A,** Z-score and % change in the RMS signal in the ipsilateral and contralateral biceps activity following stimulation of control animals (sham; CTRL_ipsi_: 97.45±6.09%, CTRL_contra_: 92.33±6.17, two-Way ANOVA stim x side: Fstim(1,401)=0.18, P=0.67, Fside(1,401)=3.12, P=0.078, Finteraction(3,401)=18.30, P=0.093). **B-C,** Response latency (two-way ANOVA target x side:F_target_(1,206)=14.24, P=0.0002, F_side_(1,206)=0.14, P=0.71, F_interaction_(1,206)=0.05, P=0.82) and response duration (two-way ANOVA target x side:F_target_(1,212)=371.60, P=0.00001, F_side_(1,212)=0.52, P=0.47, F_interaction_(1,212)=1.39, P=0.23) of the change in muscle activity following stimulation of PPN or CnF glutamatergic neurons (ipsilateral vs contralateral biceps).

**Table 1. Abbreviation of major structures reported in the manuscript.**

**Table 2. Morphological and functional parameters of PPN and CnF glutamateric neurons. Related to Figure 4.**

**Table 3. Percentages of functional subtypes of PPN and CnF glutamatergic neurons. Related to Figure 4.**

**Video 1. Frequency-dependent modulation of locomotion in the CnF. Related to Figure 5 and Supplementary Figure 4.** Example of a VGLUT2-cre mice injected in the CnF with AAV-DIO-ChR2-YFP and tested in a large open field using a progressive stimulation protocol ranging from 0.1Hz to 30 Hz.

**Video 2. Modulation of gait by PPN neurons. Related to Figure 6 and Supplementary Figure 4.** Example of a VGLUT2-cre mice injected in the PPN with AAV-DIO-ChR2-YFP and optogenetically stimulated during the elevated grid walk test.

**Video 3. Activation of locomotor muscles following CnF stimulation. Related to Figure 7 and Supplementary Figure 5**. Example of a VGLUT2-cre mice injected in the CnF with AAV-DIO-ChR2-YFP receiving optogenetic stimulation (1 and 5Hz) while held in a tail-lifted position.

