## Supplementary Figures for "Modulation of motor behavior by the mesencephalic locomotor region"

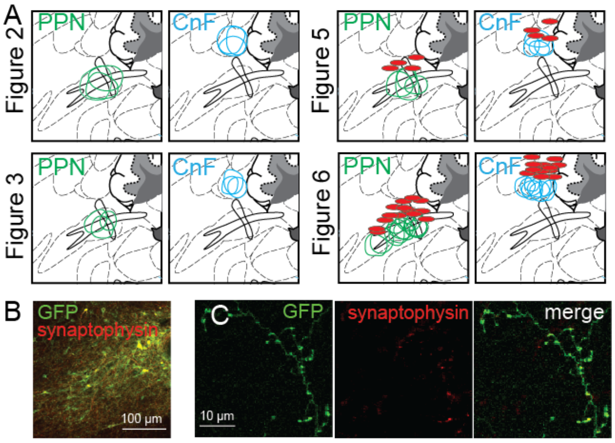


**Supplementary Figure 1.Histological analysis. Related to Figures 2, 3, 5 and 6**.
**A,** Virus spread (circles) and locations of the tip of the optic fibers (red circle) for PPN (green) and CnF (blue) groups. **B,** Fluorescent micrographs of PPN glutamatergic neurons expressing GFP and synaptophysin. **C,** High-resolution images of a transduced CnF glutamatergic axon in PAG expressing GFP in the shaft and boutons and synaptophysin-mRuby in the terminals.

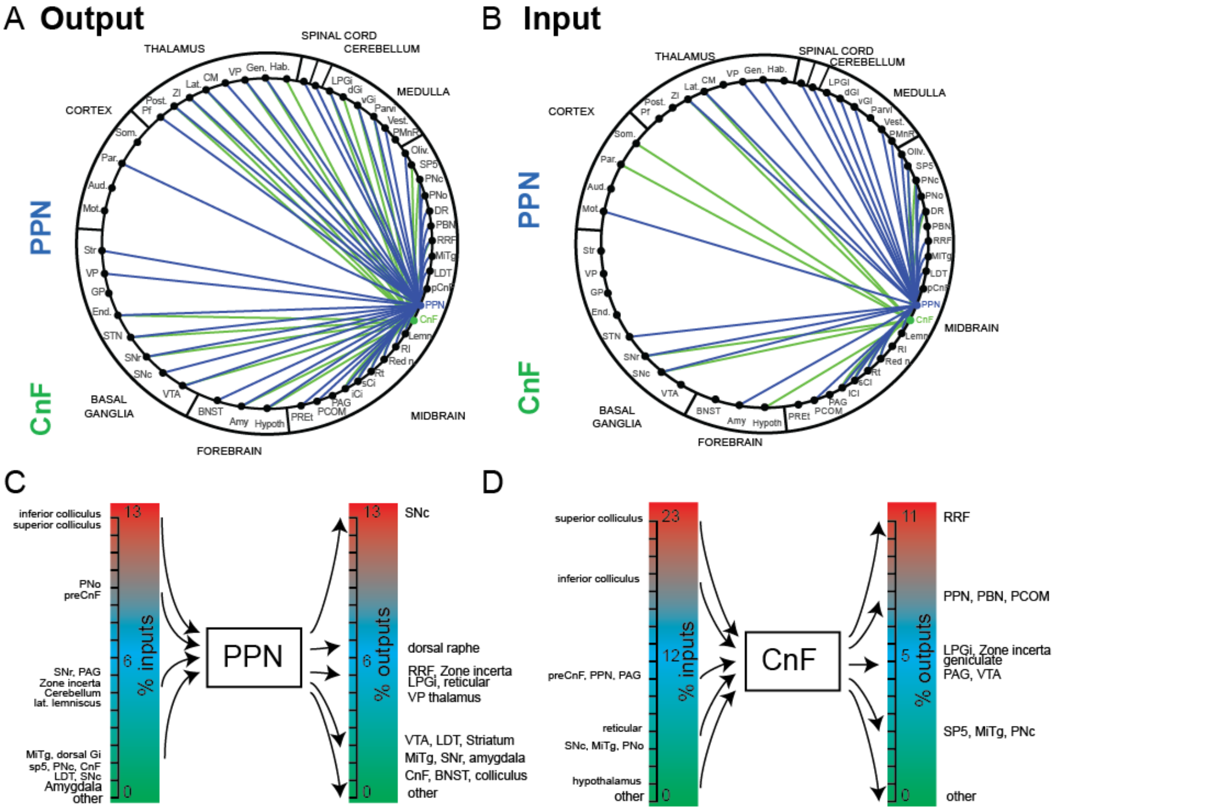


**Supplementary Figure 2. Input/output relationship of PPN and CnF**. **Related to Figures 2 and 3**.
**A**, Schematic summary of PPN (blue) and CnF glutamatergic (green) axonal distribution using relative synaptic density. **B**, Schematic summary of PPN (blue) and CnF glutamatergic (green) neuron inputs. **C-D,** Graphical representation of inputs and outputs of PPN and CnF glutamatergic neurons based on the data presented in Figures 2 and 3. Input arrows (left) are defined based on the normalized distribution of inputs neurons. Output arrows (right) are defined based on the normalized distribution of the synapses.

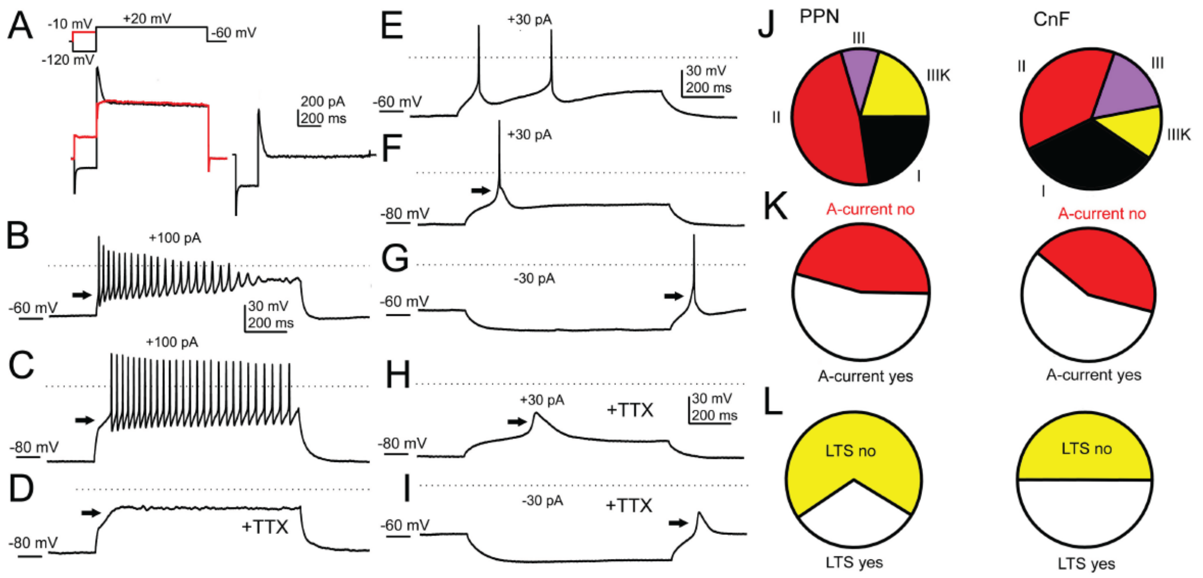


**Supplementary Figure 3. Membrane properties of the PPN and CnF glutamatergic neurons. Related to Figure 4.**

**A,** A-current was observed on most PPN and CnF glutamatergic neurons. Current traces elicited by +20 mV voltage step, preceded by -120 mV (black) and -10 mV (red) voltage steps (example shows PPN). The left current trace is the difference of the black and red current traces. **B-D,** Representative examples of the firing properties of PPN glutamatergic neurons. Trains of action potentials elicited by 100 pA depolarizing current injection from -59 mV (**B**), -87 mV (**C**) and -73 mV resting membrane potential (**D**, in the presence of TTX; the arrow indicates the lack of delay). **E-I,** Depolarization and action potential firing elicited by 30 pA depolarizing square current injection from -66 mV (**E**) and -83 mV (**F**) resting membrane potentials. Note the low threshold depolarizing spike (black arrow). (**G**) 30 pA hyperpolarizing current injection from -53 mV resting membrane potential revealed rebound spike and firing (black arrow). **H-I,** Low threshold spike and rebound depolarizing spike (respectively; arrow) in the presence of TTX. **J,** Distributions of functional neuronal types in the PPN and CnF. Group I neurons display low threshold depolarizing spikes but lack A-current (PPN: 22.7%, CnF 33.3%). Group II neurons display A-current (PPN: 47.7%, CnF 37.5%). Group III neurons display both (PPN: 9.1%, CnF 16.6%). Group IIIK lacks all (PPN: 20.5%, CnF 12.5%). **K,** Proportion of PPN and CnF neurons displaying A-current. **L,** Proportion of PPN and CnF neurons displaying low threshold spikes (LTS). All experiments have been replicated at least 3 times. All data are represented as mean ± SEM.

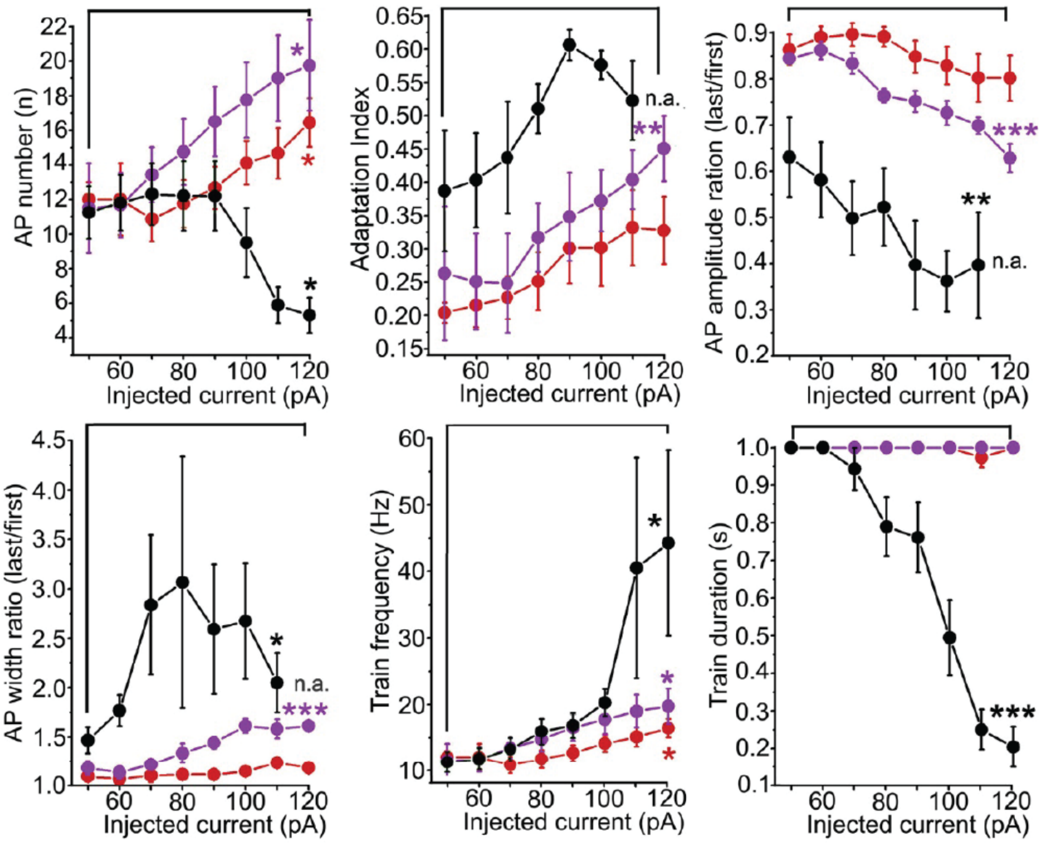


**Supplementary Figure 4. Physiological properties of MLR glutamatergic neurons. Related to Figure 4.**

Statistical summary of the number of action potentials elicited by 1-s depolarizing step, the adaptation index, the ratio of the amplitude of the last and first action potentials of the train, the ratio of the width of the last and first action potentials of the train, the frequency and the duration of the train in different functional subgroups (non-adapting, red; slowly adapting, purple; rapidly adapting, black). The significance was calculated between the first and last datapoints within each trace. * $P<0.05$, **$P<0.01$, ***$P<0.001$. All experiments have been replicated at least 3 times. All data are represented as mean ± SEM.

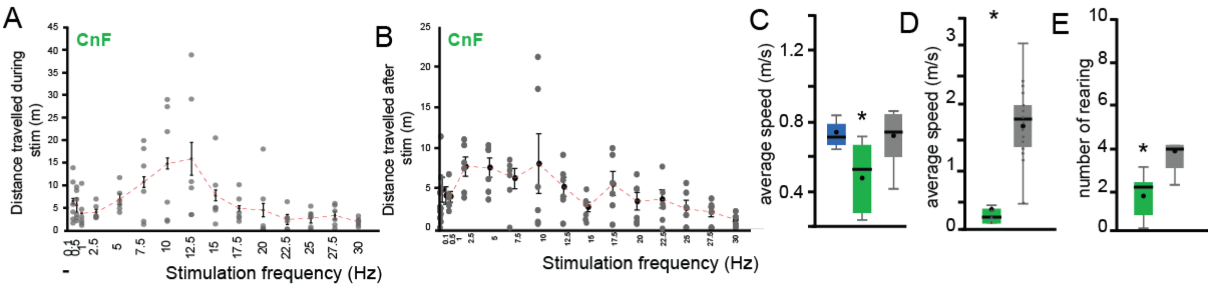


**Supplementary Figure 5. Frequency-dependent modulation of locomotion in the CnF. Related to Figures 5 and 6**.

**A-B,** Distance traveled following optogenetic stimulation (1s ON/9s OFF) during and immediately after optogenetic stimulation of CnF glutamatergic neurons using a randomized stimulation protocol ranging from 0.1Hz to 30 Hz (mixed ANOVA; *during*: F(14,89)=2.69, P=0.003, trendline: $R^2$ = 0.4774, Max: 12.5, Posthoc Bonferroni P=0.034; *after*: F(14,89)=1.24, P=0.2994). Gray dots represent individual data points, black dots represents average values, vertical lines represent SEM, and the red line represents the best fitted trendline ($y=-0.0314x^2+0.7389x+4.9582$). **C,** Average speed of animals in the CnF (blue), PPN (green) and control groups (gray) during the treadmill test (F(2,21)=17.41, P=0.0001, Bonferonni posthoc $P_{PPN\_CTRL}$=0.714, $P_{CTRL\_CNF}$=0.0001, $P_{PPN\_CNF}$=0.001). **D,** Average speed of mice in the PPN (green) and control groups (gray) during the elevated grid walk test (t-test two tail: t(28)=6.39, P=0.00001,). **E**, Total number of rearing events observed in the elevated grid walk test following stimulation of PPN glutamatergic neurons (green) and compared to control mice (gray; t(15)=2.63, P=0.0095).

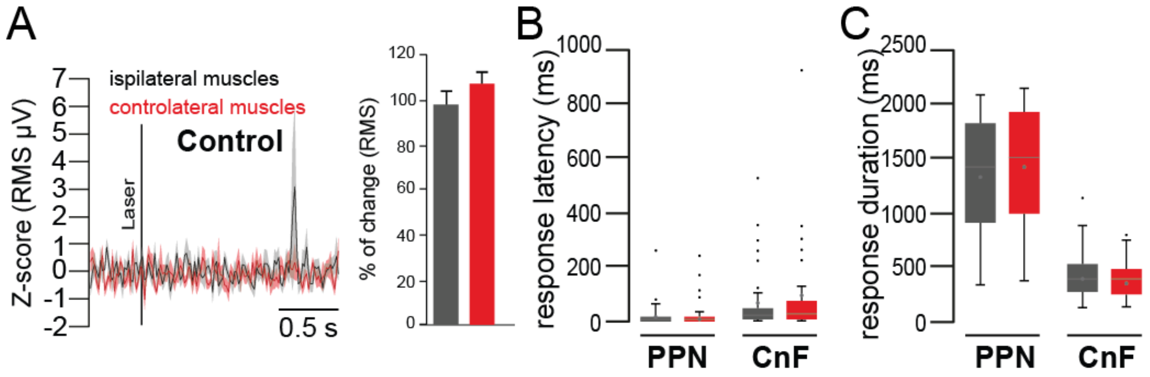


**Supplementary Figure 6. Complementary EMG experiments. Related to Figure 7**.

**A,** Z-score and % change in the RMS signal in the ipsilateral and contralateral biceps activity following stimulation of control animals (sham; $CTRL_{ipsi}$: 97.45±6.09%, $CTRL_{contra}$: 92.33±6.17, two-Way ANOVA stim x side: Fstim(1,401)=0.18, P=0.67, Fside(1,401)=3.12, P=0.078, Finteraction(3,401)=18.30, P=0.093). **B-C,** Response latency (two-way ANOVA target x side:$F_{target}$(1,206)=14.24, P=0.0002, $F_{side}$(1,206)=0.14, P=0.71, $F_{interaction}$(1,206)=0.05, P=0.82) and response duration (two-way ANOVA target x side:$F_{target}$(1,212)=371.60, P=0.00001, $F_{side}$(1,212)=0.52, P=0.47, $F_{interaction}$(1,212)=1.39, P=0.23) of the change in muscle activity following stimulation of PPN or CnF glutamatergic neurons (ipsilateral vs contralateral biceps).
