## Supplementary material for "Modulation of motor behavior by the mesencephalic locomotor region": Tables

| Abbreviation | Structure |
|---|---|
| Amy | Amygdala |
| Aud | auditory cortex |
| BNST | Bed nucleus stria terminalis |
| CM | Centromedial thalamic n. |
| DR | Dorsal raphe |
| End | Endopeduncular nucleus |
| Gen | Geniculate nucleus |
| Gi | Gigantocellular nucleus |
| GP | Globus pallidus |
| Hab | Habenula |
| Hypoth | Hypothalamic nucleus |
| iCi | Inferior colliculus |
| Lat | Lateral |
| LDT | Laterodorsal tegmental nucleus |
| Lemn | Lemniscus nucleus |
| LPGi | Lateroposterior gigantocellular nucleus |
| M1 | Primary motor cortex |
| MiTg | Microcellular tegmental nucleus |
| Mot | Motor cortex |
| n. | Nucleus |
| Oliv | Olivary nucleus |
| PAG | Periaqueductal gray nucleus |
| Parvi | Parvicellular nucleus |
| PCOM | Nucleus of the posterior commissure |
| Pf | Parafascicular thalamic nucleus |
| PMnR | Pontine mesencephalic nucleus |
| PNc | Pontine *caudalis* |
| Pno | Pontine *oralis* |
| Post. | posterior |
| preCnF | Precuneiform nucleus |
| RRF | Retrorubral field |
| Rt | Reticular nucleus |
| SC | Spinal cord |
| sCi | Superior colliculus |
| SNc | Substantia nigra *compacta* |
| SNl | Substantia nigra *lateralis* |
| SNr | Substantia nigra *reticulata* |
| Som | Somatosensory cortex |
| SP5 | Spinal trigeminal nucleus |
| STN | Subthalamic nucleus |
| STR | Striatum |
| Vest | Vestibular nucleus |
| VP | Ventral pallidum |
| VTA | Ventral tegmental area |
| ZI | Zona incerta |

**Table 1. Abbreviations.**

| **Morphological parameters** | | | |
|---|---|---|---|
| **Parameter** | **PPN** | **CnF** | **p** |
| Dendrite number | 3.66 ± 0.33 | 5.11 ± 0.42 | **0.007** |
| Node number | 8.08 ± 1.51 | 15 ± 4.38 | **0.05** |
| Ending number | 11.58 ± 1.73 | 20 ± 4.56 | **0.03** |
| Spine number | 28.83 ± 8.7 | 32.11 ± 15.08 | **0.42** |
| ChAT positivity (%) | 3.4 ± 1.8 % | 0 % | |
| n | 77 | 41 | |
| **Functional parameters** | | | |
| **Parameter** | **PPN** | **CnF** | **p** |
| A-current amplitude (pA) | 395.13 ± 0.33 | 352.49 ± 52.15 | 0.32 |
| M-current amplitude (pA; at -40 mV) | 6.38 ± 1.76 | 11.5 ± 2.41 | 0.04 |
| Persistent Na-current amplitude (pA) | 26.61 ± 5.41 | 5.31 ± 3.11 | 0.005 |
| Action potential delay at -60 mV resting membrane potential (ms) | 31.27 ± 4.07 | 27.23 ± 7.95 | 0.311 |
| Action potential delay at -80 mV resting membrane potential (ms) | 61.39 ± 8.9 (p=0.0009 between delays at -60 and -80 mV) | 44.67 ± 10.64 (p=0.09 between delays at -60 and -80 mV) | 0.169 |
| Low threshold depolarization amplitude (mV) | 17.65 ± 3.09 | 12.07 ± 2.17 | |
| Rebound depolarization amplitude (mV) | 15.17 ± 2.99 | 13.09 ± 1.79 | |
| Input resistance (MΩ) | 818.93 ± 41.94 | 792.54 ± 77.9 | 0.21 |

**Table 2. Morphological and functional parameters of PPN and CnF glutamatergic neurons. Related to Figure 4.**

| **PPN** (n = 44) | % | **CnF** (n = 24) | % |
|---|---|---|---|
| type I (LTS) | 22.7 | type I (LTS) | 33.3 |
| type II (first AP delay) | 47.7 | type II (first AP delay) | 37.5 |
| type III (LTS and first AP delay) | 9.1 | type III (LTS and first AP delay) | 16.6 |
| type IIIK (none of above) | 20.5 | type IIIK (none of above) | 12.5 |

**Table 3. Percentages of functional subtypes of PPN and CnF glutamatergic neurons. Related to Figure 4.**
